# Post-transcriptional Modifications of the Large Ribosome Subunit Assembly Intermediates in *E. coli* Expressing Helicase-Inactive DbpA Variant

**DOI:** 10.1101/2025.02.04.636506

**Authors:** Luis A. Gracia Mazuca, Jonathon E. Mohl, Samuel S. Cho, Eda Koculi

## Abstract

RNA post-transcriptional modifications are ubiquitous across all organisms and serve as fundamental regulators of cellular homeostasis, growth, and stress adaptation. Techniques for the simultaneous detection of multiple RNA modifications in a high-throughput, single-nucleotide-resolution manner are largely absent in the field, and developing such techniques is of paramount importance. We used the *Escherichia coli* ribosome as a model system to develop novel techniques for RNA post-transcriptional modification detection, leveraging its extensive and diverse array of modifications. For modification detection, we performed reverse transcriptase reactions in the presence of Mn^2^⁺ and quantified the reverse transcriptase deletions and misincorporations at modification positions using Illumina next-generation sequencing. We simultaneously detected the following modifications in ribosomal RNA (rRNA): 1-methylguanosine (m^1^G), 2-methylguanosine (m^2^G), 3-methylpseudouridine, N^6^,N^6^-dimethyladenosine, and 3-methyluridine, without chemical treatment. Furthermore, subjecting the rRNA samples to 1-cyclohexyl-3-(2-morpholinoethyl) carbodiimide metho-*p*-toluenesulfonate followed by alkaline conditions allowed us to simultaneously detect pseudouridine, 7-methylguanosine (m^7^G), 5-hydroxycytidine (OH^5^C), 2-methyladenosine, and dihydrouridine (D). Finally, subjecting the rRNA samples to KMnO_4_ followed by alkaline conditions allowed us to simultaneously detect m^7^G, OH^5^C, and D. Our results reveal that m^1^G, m^2^G, m^7^G, and D are incorporated prior to the accumulation of the 27S, 35S, and 45S large subunit intermediates in cells expressing the helicase-inactive R331A DbpA construct. These intermediates belong to three distinct stages and pathways of large subunit ribosome assembly. Therefore, our results identify the time points in three pathways at which m^1^G, m^2^G, m^7^G, and D are incorporated into the large ribosome subunit and provide a framework for broader studies on RNA modification dynamics.

## INTRODUCTION

RNA post-transcriptional modifications are ubiquitous across all kingdoms of life and play fundamental roles in regulating RNA structure, RNA–protein interactions, RNA transport, and decay ^1–7^. Furthermore, these modifications are dynamically altered in response to environmental and cellular changes ^1, 8–14^. Consequently, due to their critical roles in regulating cellular functions under varying conditions, RNA modifications influence organisms’ growth, proliferation, stress adaptation, and survival ^1^. To date, approximately 170 distinct RNA modifications have been identified ^15^. However, high-throughput, single-nucleotide-resolution techniques exist for only a limited subset of these modifications ^16^. Moreover, most of these techniques are designed to detect a single modification at a time, while detecting multiple modifications requires diverse experimental workflows with distinct bioinformatics pipelines, which are time-consuming, labor-intensive, and expensive ^16–18^. Since RNA modification roles have been shown to be interconnected, straightforward, high-resolution, and high-throughput techniques capable of detecting multiple RNA modifications simultaneously are essential for deciphering how they regulate cellular functions ^2^. In this study, we introduce novel techniques capable of simultaneously detecting different classes of RNA modifications at single-nucleotide resolution and in a high-throughput manner. The *Escherichia coli* (*E. coli*) ribosome was used as a model system due to its extensive and diverse modification content.

The *E. coli* ribosome (70S) is an RNA–protein macromolecule composed of the small (30S) and large (50S) subunits, both of which contain a plethora of RNA modifications^19^. The large subunit contains two ribosomal RNA (rRNA) molecules: 23S and 5S ^19^. The 5S rRNA, which is 120 nucleotides long, has no RNA modifications ^19, 20^. In contrast, the 23S rRNA, which is 2907 nucleotides long, contains the following RNA modifications (Table S1): 1-methylguanosine (m^1^G) 747, pseudouridine (Ψ) 748, 5-methyluridine (m^5^U) 749, Ψ 957, 6-methyladenosine (m^6^A) 1620, 2-methylguanosine (m^2^G) 1837, Ψ 1915, 3-methyl-Ψ (m^3^Ψ) 1919, Ψ 1921, m^5^U 1943, 5-methylcytidine (m^5^C) 1966, m^6^A 2034, 7-methylguanosine (m^7^G) 2073, 2’-O-methylguanosine (G_m_) 2255, m^2^G 2449, dihydrouridine (D) 2453, Ψ 2461, 2’-O-methylcytidine (C_m_) 2502, 5-hydroxycytidine (OH^5^C) 2505, 2-methyladenosine (m^2^A) 2507, Ψ 2508, 2’-O-methyluridine (U_m_) 2556, Ψ 2584, Ψ 2608, and Ψ 2609 ^19, 21–31^. The small subunit contains a single RNA molecule, the 16S rRNA, which is 1541 nucleotides long and contains the following modifications (Table S1): Ψ 516, m^7^G 527, m^2^G 996, m^5^C 967, m^2^G 1207, N4,2’-O-dimethylcytidine (m^4^C_m_) 1402, m^5^C 1407, 3-methyluridine (m^3^U) 1498, m^2^G 1516, N⁶,N⁶-dimethyladenosine (m^6^ A) 1518, and m^6^ A 1519 ^19, 23, 32–37^. With few exceptions, unique and specialized enzymes incorporate each modification into the bacterial ribosome (Table S1). In some cases, these modification enzymes function as RNA chaperones, and the modifications themselves regulate ribosome assembly ^2, 6, 7, 38^. Therefore, for a complete understanding of the ribosome maturation process, which may vary significantly under different cellular and environmental conditions, it is essential to determine the extents and roles of RNA modifications at each stage of ribosome assembly and within each pathway, as well as the specific points during the maturation process at which modification enzymes act along distinct pathways ^2, 8, 14^.

In this study, the m^1^G, m^2^G, m^3^Ψ, m^3^U, and m^6^ A modifications are simultaneously detected in rRNA without requiring chemical treatment. These modifications were previously detected either individually or in pairs using Illumina next generation sequencing (NGS) by counting reverse transcriptase stops, deletions, and/or misincorporations (Table S2). However, methods for the simultaneous detection of these modifications in a high-throughput manner with single-nucleotide resolution have been absent in the field (Table S2) ^16^. Furthermore, methods for the simultaneous detection of m^2^A, OH^5^C, m^7^G, Ψ, and D modifications have also been absent in the field (Table S2). In this study, however, the m^2^A, OH^5^C, m^7^G, Ψ, and D modifications are simultaneously detected by treating rRNA samples with 1-cyclohexyl-3-(2-morpholinoethyl) carbodiimide metho-*p*-toluenesulfonate (CMCT), followed by NaHCO₃ treatment.

In addition, we can simultaneously detect m^7^G, D, and OH^5^C modifications by subjecting rRNA to KMnO₄ plus NaHCO₃ treatment. These three modifications have been detected either individually or simultaneously in earlier studies through chemical treatment and Illumina NGS (Table S2). For example, Marchand *et al*. detected m^7^G, D, and OH^5^C by subjecting the RNA to alkaline conditions, followed by aniline chemical cleavage, and using Illumina NGS to determine the cleavage sites ^39^. However, since the aniline cleavage detects the end of the transcript, the efficacy of this powerful assay diminishes when modifications are closely spaced in sequence. By contrast, our KMnO₄ plus NaHCO₃ method, combined with Illumina NGS, counts reverse transcriptase deletions and mismatches rather than stops, making it more effective at resolving closely spaced modifications. Thus, our approach complements and extends the aniline cleavage approach by enhancing the detection of m^7^G, OH^5^C, and D modifications that are closely spaced in sequence ^39^.

We applied the Illumina NGS techniques developed in this study to measure the extents of m^1^G, m^2^G, m^7^G, and D modifications in the 27S, 35S, and 45S large subunit intermediates accumulating in cells expressing the helicase-inactive R331A DbpA construct ^40–43^. The DbpA protein belongs to the DEAD-box family of RNA helicases, which are involved in the ribosome maturation process of most known organisms ^44–51^. The extents of m^2^G, m^7^G, and D modifications in the 27S, 35S, and 45S large subunit intermediates have not previously been examined. In contrast, our previous studies determined the extents of m^2^A, OH^5^C, m^3^Ψ, and Ψ modifications in the same intermediates*_42, 43_*.

Based on the sedimentation coefficient and protein composition, the 27S, 35S, and 45S represent progressively more advanced stages of large subunit ribosome assembly, with the 27S being very early-stage, the 35S early-stage, and the 45S late-stage ^40, 41^. Pulse-labeling experiments revealed that the 27S, 35S, and 45S intermediates mature to form the 50S through independent pathways^40^. Since these intermediates represent distinct stages and pathways of large subunit ribosome assembly, our experiments identify the specific stages in these pathways at which m^1^G, m^2^G, m^7^G, and D modifications are incorporated into the 23S rRNA. Furthermore, the stages of large subunit ribosome assembly during which the D modification is incorporated into *E. coli* 23S rRNA have not been determined under any cellular or environmental conditions ^52–54^. Therefore, this is the first study to determine the specific stages during large subunit ribosome assembly at which the D modification is incorporated into *E. coli* 23S rRNA.

## MATERIALS AND METHODS

### Ribosome Particles Isolation

The 27S, 35S, and 45S large subunit intermediates, as well as the 30S and 50S subunits from wild-type cells, were isolated using the same procedure described in our previous studies ^40, 42, 43^. In brief, large subunit particles were isolated from *E. coli* BLR (DE3) pLysS Δ*dbpA*:kanR cells expressing either the wild-type DbpA or the R331A DbpA construct from the pET-3a vector ^40, 42, 43, 48, 49^. Cells were grown at 37°C in 200 mL of medium containing 10 g/L tryptone, 1 g/L yeast extract, 10 g/L NaCl, 100 μg/mL carbenicillin, and 34 μg/mL chloramphenicol. Once the cell culture reached an optical density (OD_600_) of approximately 0.3 at 600 nm, cell growth was arrested by the addition of 200 mL of ice. The cells were then pelleted by centrifugation at 6500 × g for 10 minutes at 4°C, and the supernatant was discarded. Following this, the cell pellets were resuspended in a buffer consisting of 20 mM HEPES-KOH (pH 7.5), 30 mM NH_4_Cl, 1 mM MgCl_2_, 4 mM β-mercaptoethanol (BME), and 300 μg of lysozyme. The lysozyme reaction was allowed to proceed for 30 minutes on ice. This mixture was then flash-frozen in liquid nitrogen and thawed at least three times. The freezing and thawing process was repeated multiple times to ensure complete cell lysis. Next, the DNA in the lysate was digested through the addition of 20 units of RNase-free DNase I (New England Biolabs) and incubated on ice for 90 minutes. Subsequently, the lysate was clarified by centrifugation at 17,000 × g for 30 minutes at 4°C. Lastly, to separate the ribosomal particles, approximately 80 OD_260_ of the clarified lysate was loaded directly onto a 38 mL 20–40% linear sucrose gradient or stored at −80°C for subsequent loading onto the sucrose gradient ^40, 42, 43^.

Linear 20–40% sucrose gradients were prepared in a buffer containing 20 mM HEPES-KOH (pH 7.5), 150 mM NH_4_Cl, 1 mM MgCl_2_, and 4 mM BME. Gradients were prepared using the Biocomp Gradient Master device. For particle separation, the clear cell lysate was loaded onto the linear sucrose gradient, which was then spun at 174,587 × g for 16 hours at 4°C. Upon centrifugation, the gradient was fractionated using the Teledyne R1 fraction collector or the Biocomp Gradient Station ^40, 42, 43^. The fractions containing the desired ribosomal particles were pooled and used for NGS experiments or flash-frozen in small aliquots and stored at −80°C for later use.

In the 20–40% sucrose gradient employed to separate the ribosomal particles from cells expressing the R331A DbpA construct, the 45S intermediate migrates significantly differently from the 30S, 27S, 35S, and 50S particles ^40, 42, 43^. On the other hand, the 27S and 35S intermediates travel near and under the 30S small subunit peak ^40, 42, 43^. Thus, the 27S and 35S samples also contain native 30S small subunits ^40, 42, 43^. As in our previous studies, for the experiments aimed at determining the 27S, 35S intermediates’ modifications, the NGS reads were aligned to the 23S rRNA *rrlB* gene reference sequence, and the 16S rRNA reads were disregarded ^42, 43^.

The DbpA protein interacts tightly and specifically with the 50S large subunit, and the expression of R331A does not affect the native 30S small subunit ^40, 41^. Thus, we used the 16S rRNA present in the 35S large subunit intermediate to interrogate the content of 16S rRNA modifications for all the experiments in this manuscript. The only exceptions were the rRNA samples subjected to KMnO_4_, and KMnO_4_ plus NaHCO_3_. For those latter experiments, the 16S rRNA was obtained from the 30S of cells expressing wild-type DbpA protein ^42^.

### Treatment of rRNA with CMCT and NaHCO_3_, and NaHCO_3_

Chemical treatment with CMCT followed by NaHCO_3_ of the large subunit intermediates was performed as described previously for the 27S, 35S, and 45S intermediates, as well as the 30S and 50S subunits ^42, 43^. In summary, the ribosomal proteins were stripped from 23S rRNA particles by employing phenol/chloroform extraction. Naked rRNA was then treated with 170 mM CMCT for 30 minutes at 37°C in a buffer containing 50 mM bicine (pH 8.3), 7 M urea, and 4 mM EDTA in a total reaction volume of 120 µL. This reaction was stopped by adding 0.3 M sodium acetate (pH 5.5), and rRNA was subsequently ethanol-precipitated.

The CMCT–modified rRNA was incubated with 10 mM NaHCO_3_ (pH 10.4) at 37°C for 2.5 hours and at 65°C for 30 minutes. The NaHCO_3_ reaction was stopped with 0.3 M sodium acetate (pH 5.5). rRNA was again ethanol-precipitated and stored for later use in Illumina NGS library preparation. The rRNA samples treated only with NaHCO_3_ were subjected to the same procedure as the rRNA samples treated with both CMCT and NaHCO_3_, with the sole exception that these samples were not exposed to CMCT ^42, 43^.

### Treatment of rRNA with KMnO_4_ alone, and with KMnO_4_ and NaHCO_3_

Small and large subunit particles used for KMnO_4_ treatment were isolated from wild-type-DbpA - expressing cells. Ribosomal proteins were removed from the particles using phenol/chloroform extraction and the rRNA was desalted and concentrated using ethanol–precipitation. Approximately 5 µg of rRNA was subjected to 0.12 mM KMnO_4_ plus 30 mM sodium acetate (pH 4.4) in a total volume of 25 µL ^42^. The KMnO_4_ modification reaction was allowed to proceed for three or six minutes before arresting it with 2 µL of 14.3 M BME. Prior to Illumina NGS library preparation or NaHCO_3_ treatment, rRNA was concentrated and desalted by ethanol-precipitation*_42_*.

Half of 16S and 23S rRNA samples subjected to KMnO_4_ for three or six minutes and desalted by ethanol-precipitation were additionally subjected to NaHCO_3_ (pH 10.4) at 37°C for 2.5 hours, and then at 65°C for 20 minutes. The NaHCO_3_ reaction was arrested by adding 0.3 M sodium acetate (pH 5.5). The rRNA was ethanol precipitated and subsequently used for Illumina NGS library preparation.

### Illumina NGS library preparation

The Illumina NGS library was prepared as previously described for the randomer workflow of selective 2’-hydroxyl acylation and analyzed by extension of primers and mutational profiling protocol (SHAPE-MaP) ^55^. In brief, the reverse transcriptase reaction of untreated or chemically treated rRNA was performed using the SuperScript II enzyme in the presence of 6 mM Mn^2+^ and random DNA primers ^55^. The conversion of single-stranded DNA into double-stranded DNA was performed using the NEB second strand synthesis kit. The Nextera XT DNA library preparation kit was used to tag the Illumina sequencing adaptors ^55^. The libraries were multiplexed and sequenced using Illumina 2×150 paired-end MiSeq or 2×100 paired-end NovaSeq 6000 platforms.

### Data analysis

The Bowtie 2 aligner, as part of the ShapeMapper v1.2 program, was used to align the 23S rRNA NGS reads of the 27S, 35S, 45S, and 50S samples to the 23S *rrlB* gene ^55^. Furthermore, Bowtie 2 was used to align the 16S rRNA NGS reads to the *rrsB* gene ^55^. The 16S rRNA NGS reads were obtained from the 30S small subunit isolated from cells expressing wild-type DbpA, or the 30S subunit mixed with 35S intermediates. The Bowtie 2 aligner has been proven effective in aligning mixtures of NGS reads to the correct reference sequences, particularly when the RNA sequences in the mixture are sufficiently different ^56^. Given that 16S and 23S rRNA sequences do not have significant similarities, Bowtie 2 can accurately align NGS reads from a mixture of 16S and 23S rRNA, such as in the 27S and 35S samples, to their correct reference sequences ^43^.

The mutation rates for each nucleotide in all ribosomal particles were calculated as in our previous studies ^42, 43^. Briefly, the suggested parameters for ShapeMapper v1.2 for the randomer primer workflow and the Nextera XT kit were used ^55^. ShapeMapper v1.2 calculates mutation rates of a given nucleotide as the ratio of the sum of misincorporations and deletions divided by the total read depth at that nucleotide position ^55^. Thus, the mutation rate represents the fraction of reverse transcriptase reaction mismatches and deletions at a nucleotide position when compared to the total number of nucleotide reads at the same position ^55^.

### Detection of m^1^G, m^2^G, m^3^Ψ, and m^3^U post-transcriptional modifications in the 27S, 35S, 45S, 50S, and 30S ribosomal particles without chemical treatment

m^1^G, m^2^G, m^3^U, and m^3^Ψ are detected as the averages of mutation rates from two chemically untreated biological samples.

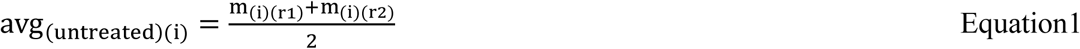

In Equation 1, avg_(untreated)(i)_ represents the average mutation rate in chemically untreated samples for nucleotide i. m_(*i*)(*r*1)_ and m_(*i*)(*r*2)_ denote the mutation rates for nucleotide i in biological replicates 1 and 2, respectively.

The standard deviations for the mutation rates of chemically untreated samples were calculated using the following equation:

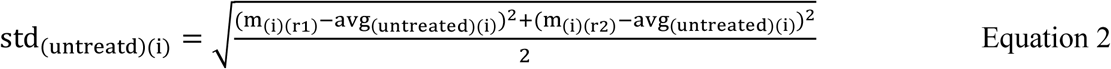

In Equation 2, std_(untreated)(i)_ represents the standard deviation of the mutation rate averages calculated with Equation 1. avg_(untreated)(i)_ denotes the average mutation rate calculated with Equation 1. m_(i)(r1)_ and m_(i)(r2)_ represent the mutation rates for nucleotide i in biological replicates 1 and 2, respectively.

### Detection of D and m^7^G post-transcriptional modifications in the 27S, 35S, 45S, 50S, and 30S ribosomal particles

D and m^7^G were detected by analyzing the mutation rate as calculated by ShapeMapper v1.2 for the rRNA samples subjected to the following alkaline conditions: NaHCO_3_, CMCT plus NaHCO_3_, and KMnO_4_ plus NaHCO_3_.

### Detection of D and m^7^G from rRNA samples treated with NaHCO_3_ alone, and with KMnO_4_ plus NaHCO_3_

NGS data were obtained from a single biological replicate for the rRNA samples subjected to NaHCO₃, KMnO₄, and KMnO₄ plus NaHCO₃ treatments. The mutation background corrections and error propagations for these experiments were performed as follows. First, the ShapeMapper v1.2 mutation rate was background-corrected by subtracting the average mutation rate of the chemically untreated sample (Equation 1) from that of the chemically treated rRNA sample.

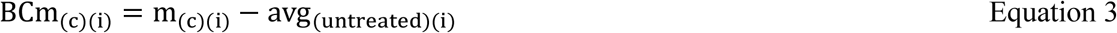

In Equation 3, BCm_(c)(i)_ represents the background-corrected mutation rate for nucleotide i of the rRNA sample subjected to one of the following chemical treatments: NaHCO_3_, KMnO_4_, and KMnO_4_ plus NaHCO_3_. m_(c)(i)_ denotes the mutation rate of nucleotide i subjected to NaHCO_3_, KMnO_4_, or KMnO_4_ plus NaHCO_3_. avg_(untreated)(i)_ denotes the average mutation rate of nucleotide i for the untreated rRNA samples, as calculated from Equation 1.

The error propagation for BCm_(c)(i)_ was performed by taking into account the errors of both m_(c)(i)_ and avg_(untreated)(i)_. The error for m_(c)(i)_ was calculated as previously described using the following equation ^57^.

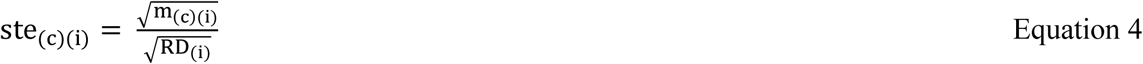

In Equation 4, ste_(c)(i)_ represents the mutation rate standard error for nucleotide i, m_(c)(i)_ denotes the mutation rate at nucleotide i, and RD_(i)_ is the total read depth at nucleotide i ^57^. The final error for BCm_(c)(i)_ was calculated through error propagation as follows.

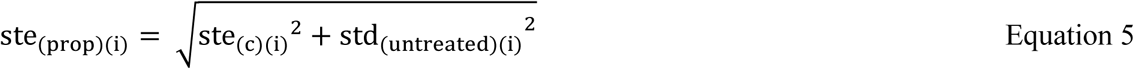

In Equation 5, ste_(prop)(i)_ represents the propagated error for BCm_(c)(i)_ when only one biological replicate was investigated. std_(untreated)(i)_ and ste_(c)(i)_ denote the errors as calculated from Equations 2 and 4, respectively.

*Detection of D and m^7^G from rRNA samples treated with CMCT plus NaHCO_3_* The ShapeMapper v1.2 mutation rates of rRNA samples subjected to CMCT plus NaHCO_3_ were background-corrected by subtracting the average mutation rates of the chemically untreated samples (Equation 1) from those of the CMCT plus NaHCO_3_-treated rRNA samples.

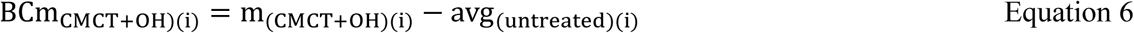

In Equation 6, m_(CMCT+OH)(i)_ represents the mutation rate of the rRNA samples subjected to CMCT plus NaHCO_3_. avg_(untreated)(i)_ denotes the average mutation rate of nucleotide i for the untreated rRNA samples, as calculated from Equation 1.

The average of the two biological replicates was obtained for the rRNA samples treated with CMCT plus NaHCO_3_ using the equation below.

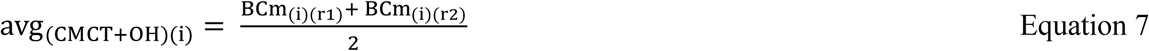

In Equation 7, avg_(CMCT+OH)(i)_ represents the average mutation rate in nucleotide i from biological replicates 1 and 2, chemically treated with CMCT plus NaHCO_3_. BCm_(i)(r1)_ and BCm_(i)(r2)_ denote the background-corrected mutation rates at nucleotide i in CMCT plus NaHCO_3_ treated biological replicates 1 and 2, as described in Equation 6.

The errors for the mutation rates of the CMCT plus NaHCO_3_-treated samples were calculated by

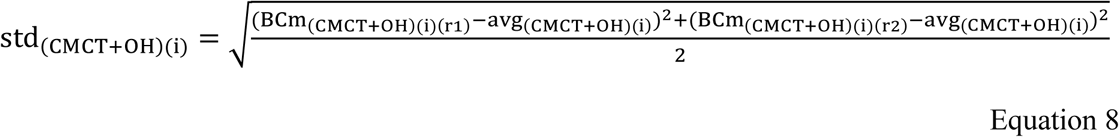

In Equation 8, std_(CMCT+OH)(i)_ represents the standard deviation from the mutation rate averages. avg_(CMCT+OH)(i)_ is the average mutation rate calculated using Equation 7. BCm_(CMCT+OH)(i)(r1)_ and BCm_(CMCT+OH)(i)(r2)_ represent the background-corrected mutation rates for nucleotide i in biological replicates 1 and 2, for samples subjected to CMCT plus NaHCO_3_, as described in Equation 6.

### Threshold used for modification detection

In our study, rRNA nucleotides were considered modified if they had a background-corrected mutation rate of 0.05 or greater in experiments with one biological replicate (Equation 3) or an average mutation rate of 0.05 or greater in experiments with two biological replicates (Equations 1 and 6).

## RESULTS AND DISCUSSION

### Detection of m^1^G, m^2^G, m^3^Ψ, m^3^U, and m^6^ A through reverse transcriptase reaction deletions and misincorporations without chemical treatment

Illumina NGS combined with ShapeMapper v1.2 was employed to detect m^1^G 747, m^2^G 1837, m^3^Ψ 1919, and m^2^G 2449 in 23S rRNA of the 50S, as well as m^2^G 1207, m^3^U 1498, and m^6^ A 1519 in 16S rRNA of the 30S (Figure 1, Table 1, and Table 2). These modifications exhibited a mutation rate equal or higher than 0.05 in both biological replicates (Figure 1, Table 1, Table 2). By contrast, the mutation rates of m^2^G 966, m^4^C_m_ 1402, m^2^G 1516, and m^6^_2_A 1518 in 16S rRNA were below our threshold of 0.05 (Figure 1B), making these modifications undetectable using this method. Importantly, employing our threshold, we observed only two false-positive modifications in both biological replicates of 23S rRNA (2907 nucleotides long) (Table S3). In 16S rRNA (1541 nucleotides long), three false-positive modifications were observed in one biological replicate and none in the other (Table S3). A strong correlation observed in two independent NGS experiments for the 23S and 16S rRNA molecules underscores the reproducibility of our modification detection method (Figure 1).

**Figure 1.**
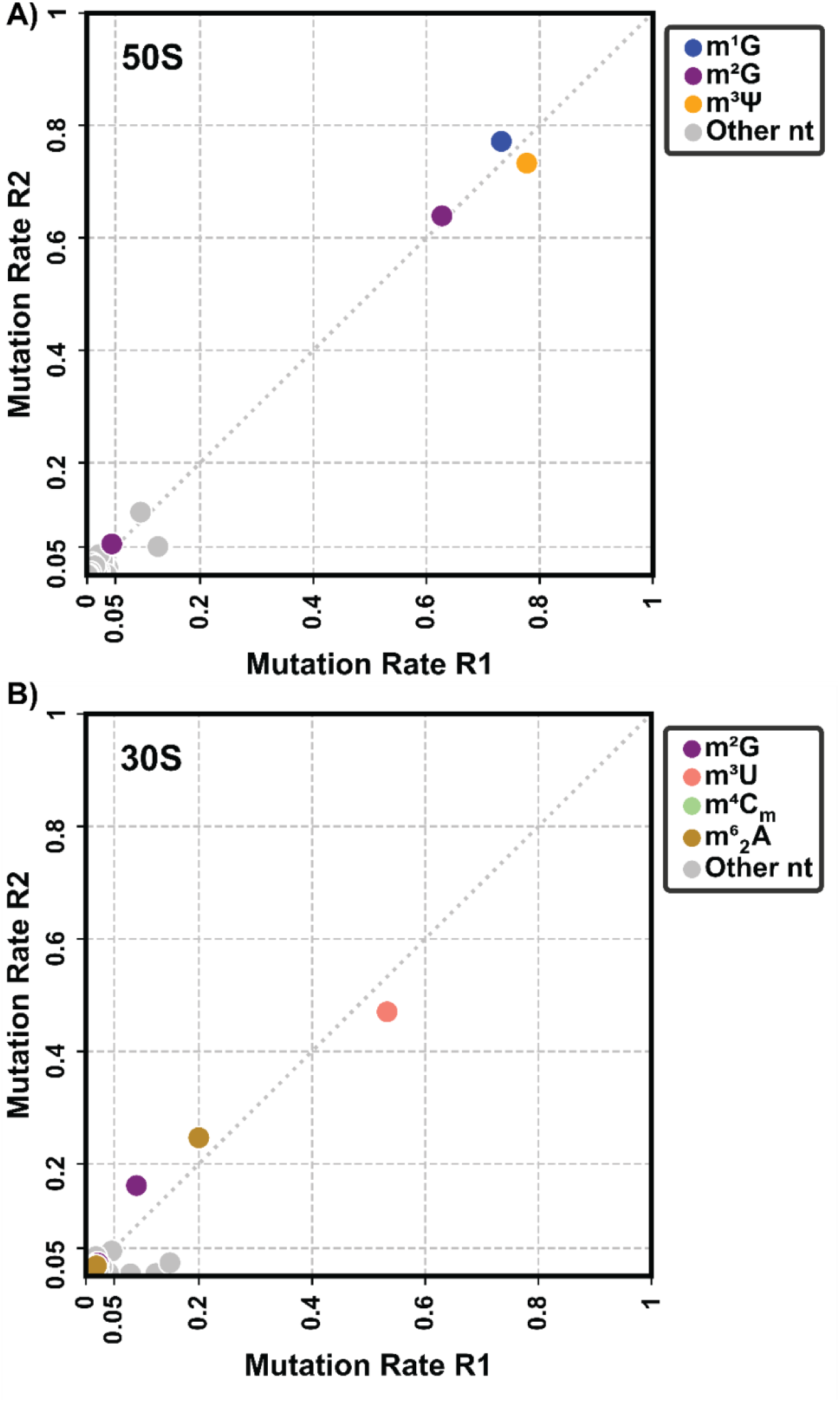
Detection of m^1^G, m^2^G, m^3^Ψ, m^3^U, and m^6^ A using reverse transcriptase reaction deletions and misincorporations. ShapeMapper v1.2 was used to count the reverse transcriptase reactions’ deletions and misincorporations. The mutation rates were calculated from deletions and misincorporations as explained in Materials and Methods. The mutation rates shown in this figure are not background-corrected. The unmodified nucleotide and the modified nucleotides that cannot be detected in these experiments are labeled as gray circles, m^1^G in blue, m^2^G in purple, m^3^Ψ in orange, m^3^U in pink, m^4^C_m_ in green, and m^6^ A in brown. R1 and R2 represent the mutation rates for the biological replicates 1 and 2. The mutation rates for the modified nucleotides agree very well between the two biological replicates. A) All the m^1^G, m^2^G, and m^3^Ψ modifications present in the 23S rRNA of mature 50S can be detected by using the threshold of 0.05 for the mutation rate (Table 1). B) The m^2^G 1207, m^3^U 1498, and m^6^ A 1519 modifications present in the 16S rRNA of mature 30S can be detected by counting reverse transcriptase reaction deletions and misincorporations and a threshold of 0.05 for the mutation rate. The m^2^G 966, m^2^G 1516, m^6^ A 1518, and m^4^C_m_ 1402 exhibit mutation rates below our threshold of 0.05; thus, they cannot be detected using our method.

**Table 1.**
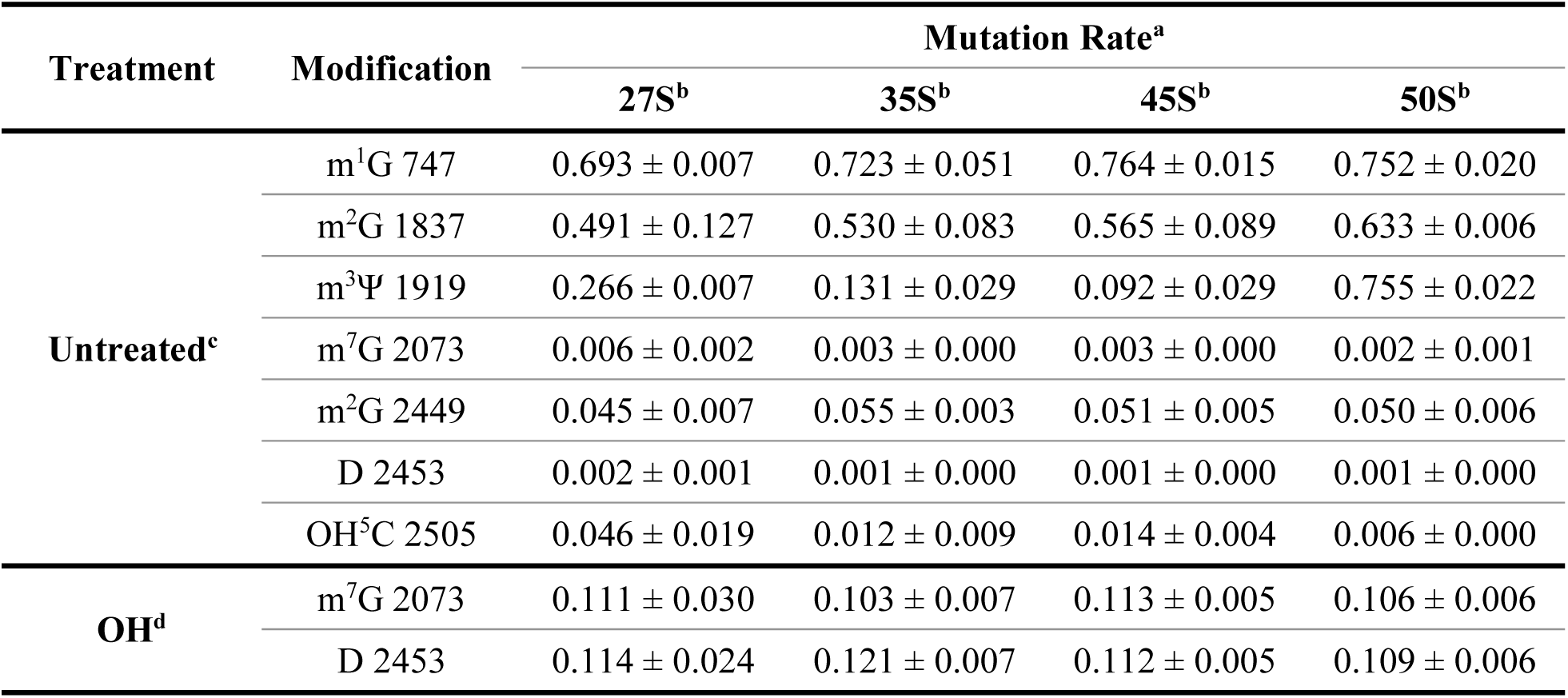
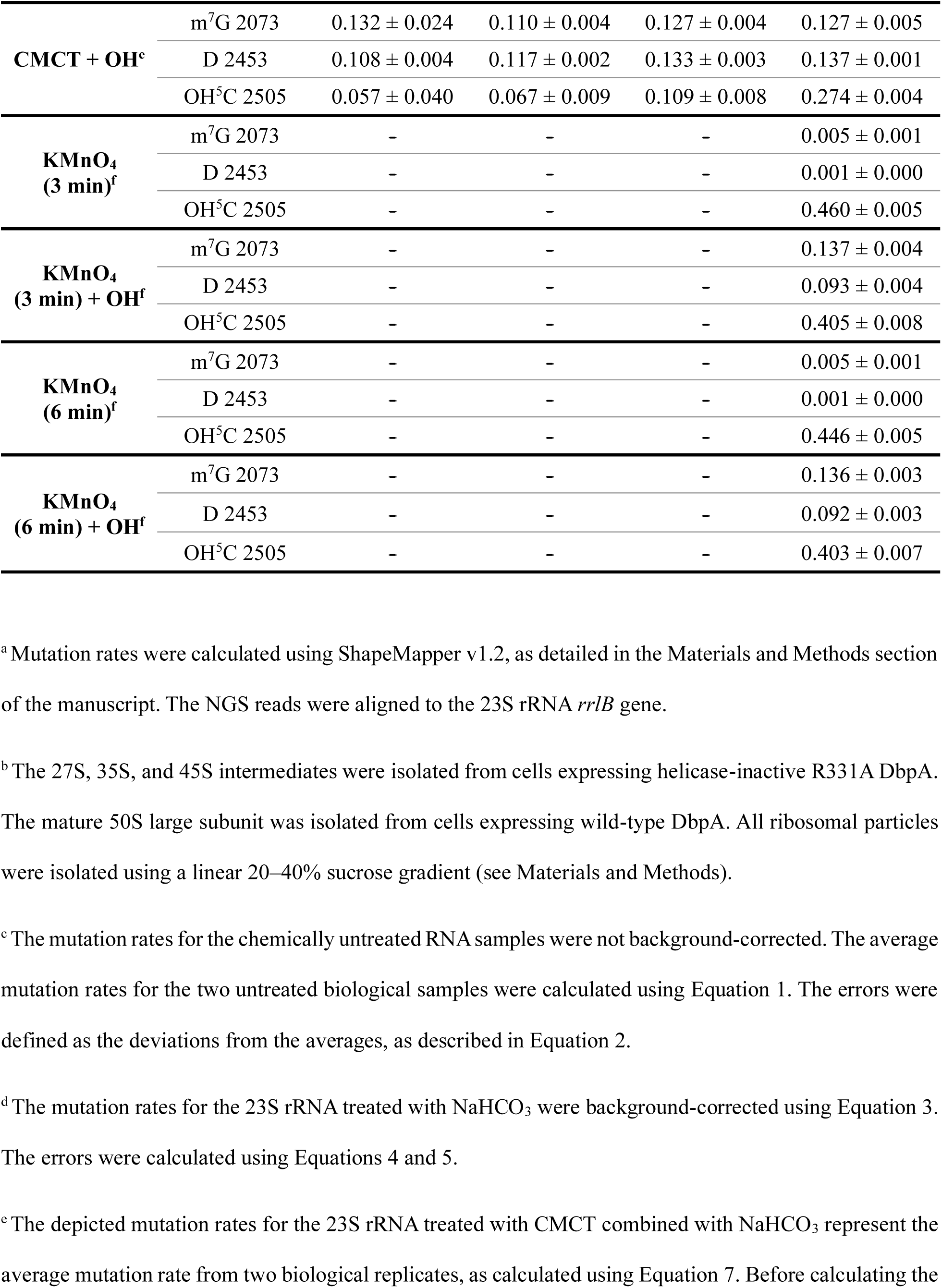
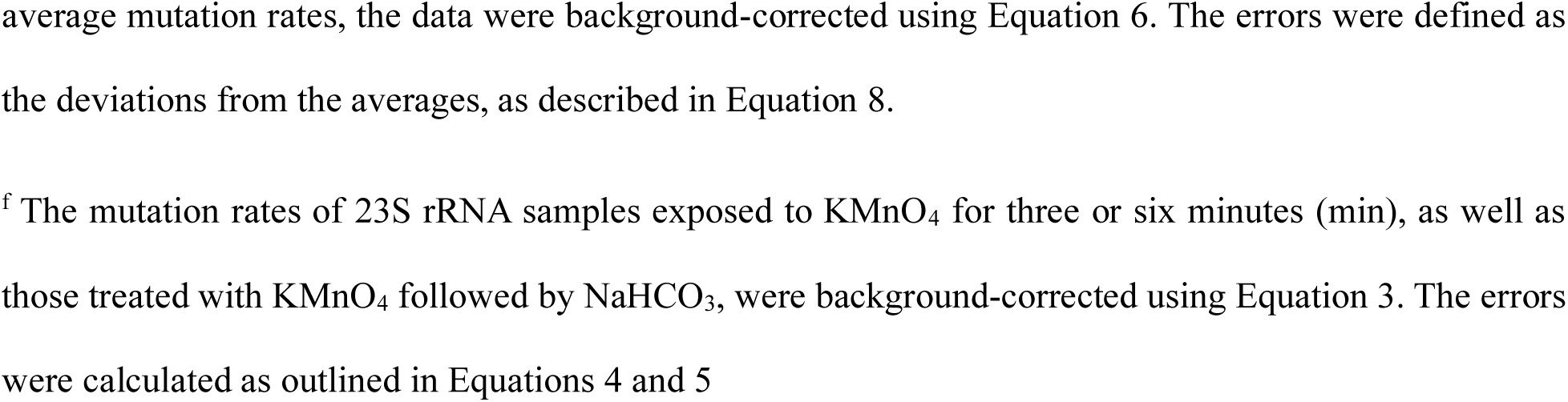
Mutation rates of m^1^G, m^2^G, m^3^Ψ, m^7^G, D, and OH^5^C modifications in 27S, 35S, and 45S intermediates and the 50S mature large subunit in the absence or presence of chemical treatments.

**Table 2.**
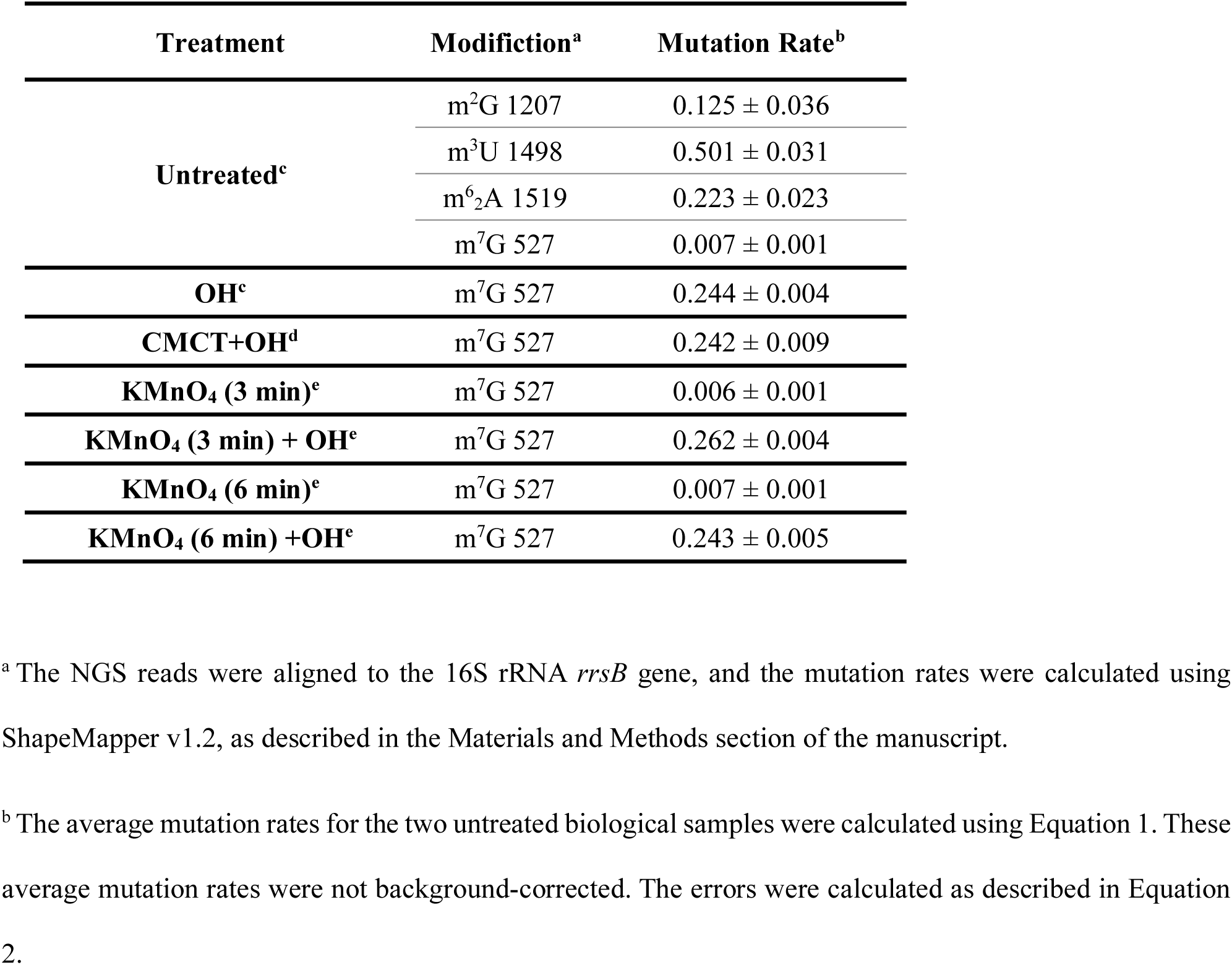
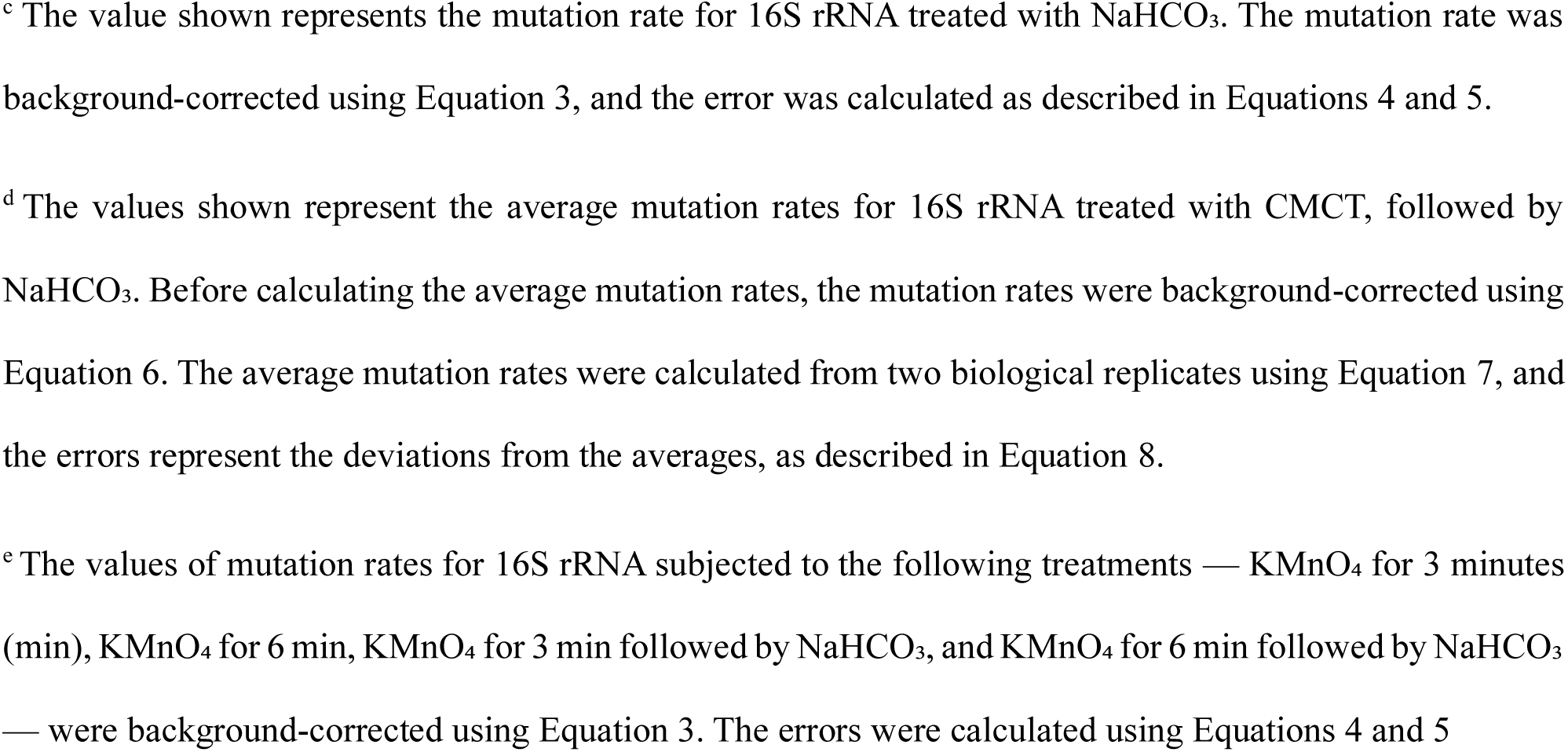
Mutation rates of m^2^G, m^3^U, m^6^_2_A, m^7^G, m^4^C_m_ in the mature 30S small ribosomal subunit in the absence or presence of chemical treatments.

The average mutation rate for the m^4^C_m_ 1402 modification in 16S is 0.017 ± 0.002, which is below the threshold mutation rate of 0.05 used in this study for modification detection, but significantly higher than the average background mutation rate of all the unmodified nucleotides of 16S rRNA, which is 0.003 ± 0.004. This suggests that our assay could potentially detect m^4^C_m_ modifications. The low mutation rate we observe could result from the inability of m^4^C_m_ modifications to induce a significant number of mismatches and deletions during the reverse transcriptase reaction. Additionally, the absolute stoichiometry of m^4^C_m_ in 16S rRNA of the mature 30S ribosome remains unknown, to the best of our knowledge. It is also possible that the m^4^C_m_ modification is incorporated at substoichiometric levels in 16S rRNA of the mature ribosome, and these low levels of incorporation may fall below the detection threshold of our assay.

Our inability to detect the m^2^G 966, m^2^G 1516, and m^6^_2_A 1518 modifications in 16S rRNA, despite successfully detecting other m^2^G and m^6^ A modifications in 16S and 23S rRNA, may result from sequence- or modification-context dependency of the reverse transcriptase reaction stops and errors (Figure 1, Table 1, and Table 2). ShapeMapper v1.2 only counts deletions and misincorporations ^55^. Thus, if certain RNA sequences or adjacent modifications increase the number of reverse transcriptase reaction stops at the m^2^G and m^6^ A positions while simultaneously reducing the mutation rates below our threshold of 0.05, these specific m^2^G and m^6^ A modifications will remain undetectable by our assay. Previous studies found that reverse transcriptase stops and misincorporation frequencies at 1-methyladenosine (m^1^A) modification sites, in chemically untreated samples, and at CMCT-modified Ψ sites are highly dependent on the sequence context ^58, 59^.

Interestingly, the patterns of mismatches and deletions for m^2^G 1837 and m^2^G 2449 in 23S rRNA and m^2^G 1207 in 16S rRNA are similar to one another and distinct from that of m^1^G 747 in 23S rRNA (Figure S1). Most often, the reverse transcriptase at the m^2^G position misincorporates a C, followed by an A, and less frequently a T (Figure S1). On the other hand, at the m^1^G 747 position, the reverse transcriptase most often skips a base, followed by misincorporating a T, and less often misincorporating an A (Figure S1). While a larger dataset needs to be analyzed with m^1^G and m^2^G modifications located in different sequence contexts, the data presented here indicate that the m^1^G and m^2^G modifications produce distinct reverse transcriptase signatures under our reaction conditions (Figure S1). These unique signatures could serve as a basis for discriminating between m^1^G and m^2^G modifications in RNA molecules with unknown modification compositions.

The m^3^Ψ 1919 in 23S rRNA and m^3^U 1498 in 16S rRNA also induce distinct reverse transcriptase signatures. At the position of m^3^Ψ 1919 in 23S rRNA, the reverse transcriptase most frequently misincorporates an A, less frequently a G, and rarely a C. In contrast, at the position of m^3^U 1498 in 16S rRNA, the reverse transcriptase most often misincorporates a G, followed by an A, and occasionally misincorporates a C or skips a base (Figure S1). Future experiments will determine whether differences in reverse transcriptase signatures occur when m^3^Ψ and m^3^U are located in different sequence contexts from those investigated here. If the distinct reverse transcriptase m^3^Ψ and m^3^U modification signatures are consistent across various RNA sequences, these signatures could provide a reliable method for distinguishing between the two modifications in an RNA molecule with unknown m^3^Ψ and m^3^U composition.

Direct RNA sequencing (DRS) using Oxford Nanopore Technologies (ONT) enables the simultaneous detection of all RNA modification classes analyzed in the manuscript *^8,^* ^14^. However, a limitation of DRS ONT in its present form is its low sequencing accuracy, which produces a significant number of false-positive modification signals ^60^. The Illumina NGS techniques described here exhibit very few false-positive modifications (Table S3). Thus, using our Illumina NGS techniques compared to DRS ONT sequencing, to discriminate between true and false modifications in an RNA molecule with an unknown modification composition, fewer mass spectrometry or other validation methods will be needed.

### Subjecting rRNA to CMCT followed by alkaline conditions enables the simultaneous detection of the m^7^G, D, m^2^A, Ψ, and OH^5^C modifications

Treatment of 23S rRNA and 16S rRNA with NaHCO₃ induces structural changes in both m^7^G and D nucleotides, leading to reverse transcriptase deletions and mismatches that exceed our mutation rate threshold of 0.05 and significantly surpass the average mutation rate of other nucleotides in both rRNA molecules (Figure 2, Table 1, and Table 2). In the NaHCO₃-treated rRNA samples, one false positive modification was observed in 23S rRNA (2907 nucleotides long) and three false positive modifications were observed in 16S rRNA (1541 nucleotides long) (Figure 2, Table S3). These false positive modifications represent a very small fraction of the total nucleotides analyzed.

**Figure 2.**
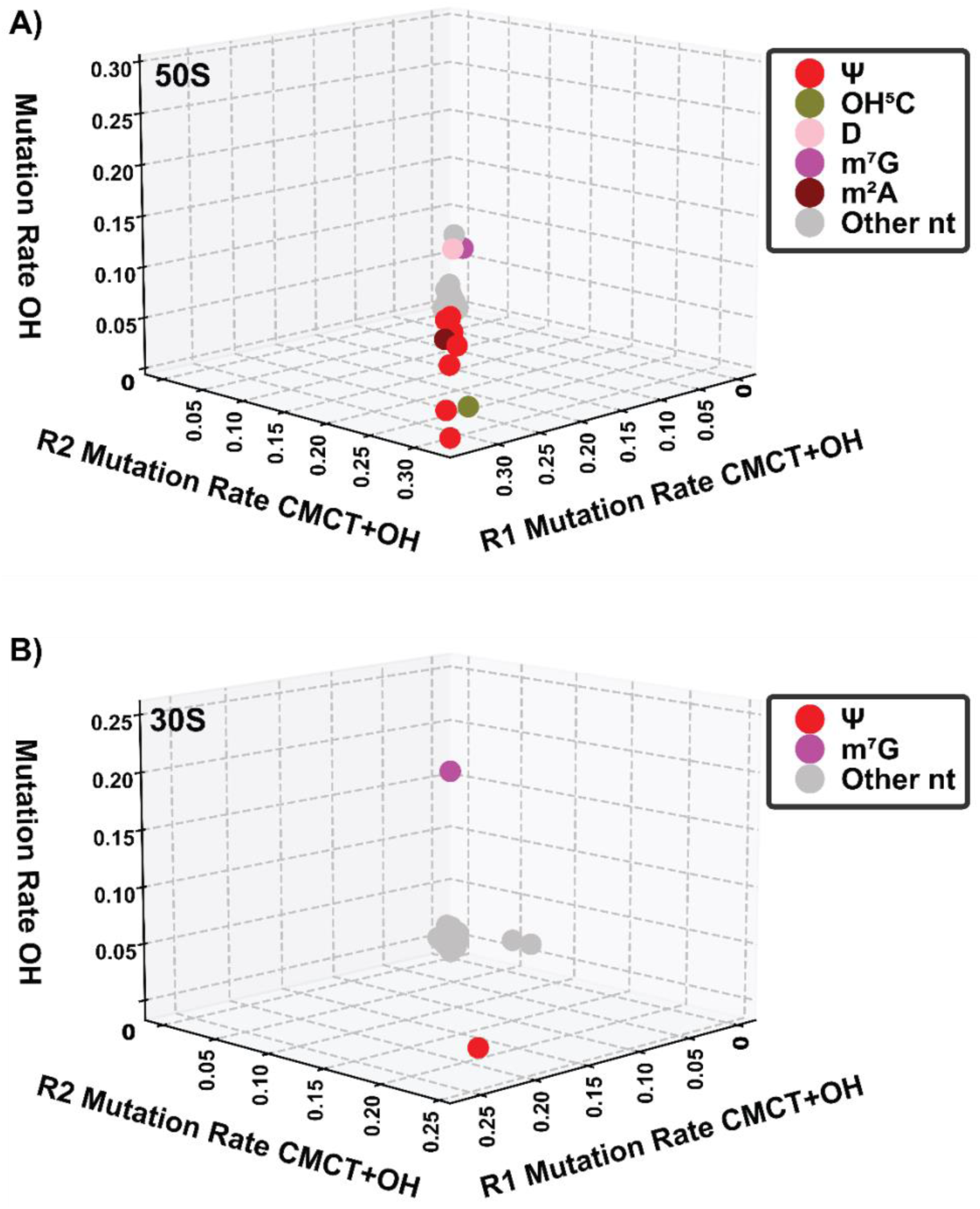
Alkaline treatment combined with CMCT enables the simultaneous detection of m^7^G, D, m^2^A, Ψ, and OH^5^C. A) 23S rRNA from the mature 50S ribosomal subunit contains one m^7^G (purple) at position 2073 and one D (pink) at position 2453 (Table S1). These modifications exhibit significantly higher mutation rates than the average mutation rate of other nucleotides in the 23S rRNA sample subjected to NaHCO₃ (y-axis). Furthermore, in the 23S rRNA samples treated with CMCT plus NaHCO₃ (x-and y-axes), the mutation rates of m^7^G at position 2073 and D at position 2453 are similar to those of Ψ modifications (red) at positions 748, 957, 1915, 1921, 2461, 2508, 2584, 2608, and 2609; OH^5^C (green) at position 2505; and m^2^A (brown) at position 2507, which are significantly higher than the average mutation rate of other nucleotides in 23S rRNA. R1 and R2 represent two biological replicates of the 50S large subunit subjected to treatment with both CMCT and NaHCO₃. B) 16S rRNA from the 30S small subunit contains one Ψ at position 516 and one m^7^G at position 527 but does not contain D or OH^5^C modifications (Table S1). Both Ψ and m^7^G modifications exhibit significantly higher mutation rates than the mutation rates of other nucleotides in 16S rRNA samples treated with CMCT plus NaHCO₃ (x-and y-axes; R1 and R2 represent two biological replicates). Additionally, the m^7^G modification exhibits a significantly higher mutation rate than the average mutation rate of other nucleotides in 16S rRNA treated only with NaHCO₃ (z-axis). These results confirm that m^7^G can be reliably detected using treatment with CMCT plus NaHCO₃ or NaHCO₃ alone. The mutation rates depicted in panels (A) and (B) have been background-corrected as described in Equations 3 and 6. In both panels (A) and (B), the abbreviation “nt” denotes nucleotide. The mutation rates for the D, m^7^G, m^2^A, OH^5^C, and Ψ modifications across the two biological replicates treated with CMCT followed by NaHCO₃ show strong agreement, demonstrating the reproducibility of our method.

Previous studies demonstrated that subjecting tRNA to alkaline conditions causes the D nucleotide to undergo ring opening, which results in reverse transcriptase stops, as identified by gel electrophoresis ^61, 62^. Under our experimental conditions, this D ring opening leads to reverse transcriptase deletions and mismatches (Figure 2, Table1, Figure S2). Additionally, exposing RNA m^7^G modifications to alkaline and/or reducing conditions has been shown to induce imidazole ring fission ^63^. Prolonged exposure to high temperatures and acidic conditions following this process can produce an abasic site ^64, 65^. However, in our experiments, we do not expose the RNA to acidic conditions at high temperatures for extended periods. Therefore, the reverse transcriptase deletions and mismatches detected in the 23S and 16S rRNA samples are likely caused by the imidazole ring fission of m^7^G in the presence of NaHCO₃ (Figure 2, Table 1, Table 2, and Figure S2).

In our previous studies, we showed that subjecting rRNA to CMCT followed by NaHCO_3_ treatment enabled the simultaneous detection of m^2^A, Ψ, and OH^5^C modifications ^42, 43^. In those studies, the mutation rates of NaHCO_3_-treated samples were used for background correction. As a result, the D and m^7^G modifications, which were altered by NaHCO_3_ in the same manner in both the NaHCO_3_-treated samples and the CMCT plus NaHCO_3_-treated samples, and which induced reverse transcriptase mismatches and deletions at similar levels, were not detected (Figure 2, Table 1, and Table 2) ^42, 43^. By using the mutation rates of chemically untreated RNA samples to perform the background correction on the mutation rates of rRNA samples treated with CMCT plus NaHCO_3_, we can simultaneously detect m^2^A, Ψ, OH^5^C, m^7^G, and D (Figure 2, Table 1, and Table 2).

Since Ψ and D are both U modifications, the CMCT plus NaHCO₃ treatment cannot discriminate between Ψ and D modifications in an RNA molecule where Ψ and D sequence positions are unknown. To address this issue, the RNA sample should also be subjected to NaHCO₃ treatment alone, which uniquely induces reverse transcriptase deletions and mismatches at D modification sites but not at Ψ modification sites (Figure 2A, Table 1). Alternatively, the pattern of reverse transcriptase deletions and misincorporations could be analyzed in RNA samples subjected to CMCT plus alkaline conditions to differentiate between D and Ψ modifications.

However, under our experimental conditions, the reverse transcriptase deletion and misincorporation patterns at Ψ modification positions in rRNA samples treated with CMCT plus NaHCO₃ were found to be sequence-context dependent (Figure S3). Moreover, the reverse transcriptase pattern of deletions and misincorporations at the only D modification site in the ribosome overlapped with the patterns observed for Ψ modifications (Figure S2, Figure S3). Thus, based on our data, D and Ψ modifications cannot be reliably differentiated using reverse transcriptase deletion and mismatch signatures alone.

The reverse transcriptase signature of mismatches and deletions differs between m^7^G 2073 of 23S rRNA and m^7^G 527 of 16S rRNA (Figure S2). This difference arises from the distinct sequence contexts where these two m^7^G modifications are located in 23S and 16S rRNA. Therefore, the reverse transcriptase deletions and mismatches observed under our experimental conditions, similar to those of Ψ modifications, are sequence-context dependent (Figure S2, Figure S3).

Previous studies demonstrated that CMCT-induced reverse transcriptase stops and misincorporations at Ψ modification sites are sequence context-dependent ^59^. Similarly, the stops and mismatches signature of reverse transcriptase at m^1^A modification sites was also found to be sequence-context dependent ^58^. Our experiments revealed that the reverse transcriptase deletions and misincorporations signatures for CMCT-modified Ψ and m^7^G under alkaline conditions are sequence-context dependent, paralleling the patterns observed in earlier studies on CMCT-modified Ψ and m^1^A modifications (Figure S2, Figure S3) ^58, 59^.

### m^7^G, D, and OH^5^C are detected simultaneously by subjecting rRNA to KMnO_4_ followed by alkaline treatment

In our previous study, we demonstrated that treating rRNA with KMnO₄ for 3 or 6 minutes alters the chemical structure of OH^5^C, resulting in reverse transcriptase mismatches and deletions ^42^. In the present study, we extended this approach by following the 3- and 6-minute KMnO₄ treatments with alkaline exposure (Figure 3, Figure S4). This sequential treatment increased the reverse transcriptase mutation rates at m^7^G, D, and OH^5^C nucleotides in the 50S rRNA and at m^7^G in the 30S rRNA, compared to other nucleotides (Figure 3). A comparison of mutation rates between rRNA samples treated with both KMnO_4_ and NaHCO_3_ and those treated with KMnO₄ alone revealed that the structural alterations at m^7^G and D nucleotides are induced specifically by the NaHCO₃ treatment, not by KMnO₄. These alterations drive the reverse transcriptase mismatches and deletions at m^7^G and D modification positions relative to other nucleotides (Figure 3, Figure S2, Figure S4, Table 1, and Table 2).

**Figure 3.**
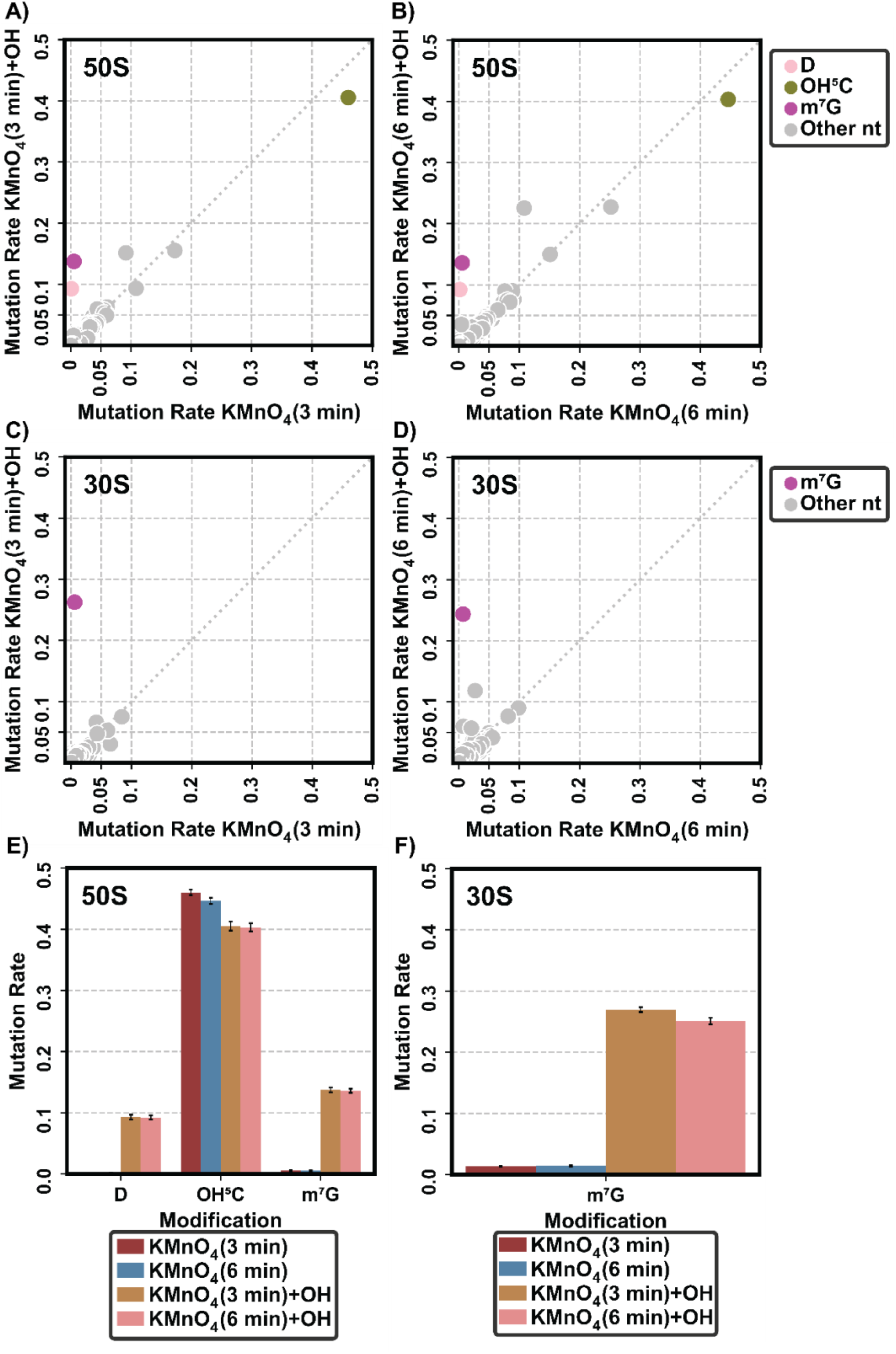
Sequential chemical treatment of rRNA with KMnO₄ followed by alkaline treatment enables simultaneous detection of m^7^G, D, and OH^5^C modifications. A) Mutation rates of m^7^G and D nucleotides in 23S rRNA treated with KMnO_4_ followed by NaHCO_3_ increase significantly when compared to 23S rRNA treated with KMnO_4_ alone. This indicates that chemical changes induced by alkaline conditions at m^7^G and D nucleotides result in an elevated reverse transcriptase mutation rate. The x-axis represents 23S rRNA treated with KMnO₄ for 3 minutes, while the y-axis represents 23S rRNA treated sequentially with KMnO₄ followed by NaHCO₃ for 3 minutes. The m^7^G (magenta), D (pink), and OH^5^C (olive green) modifications exhibit higher mutation rates in the dual-treated sample compared to the other nucleotides (gray). Consistent with our previous findings, the OH^5^C modification exhibits a higher mutation rate than that of the other nucleotides in the rRNA sample treated only with KMnO_4_ ^42^. B) The same experiment as in (A), but with 23S rRNA treated with KMnO₄ for 6 minutes instead of 3 minutes. C) In 16S rRNA, the sole m^7^G nucleotide shows a significantly elevated mutation rate when treated with KMnO₄ followed by NaHCO₃ for 3 minutes, compared to KMnO₄ alone. The x-axis represents the mutation rate for 16S rRNA treated with KMnO₄ alone, and the y-axis represents the mutation rate for 16S rRNA treated with KMnO₄ followed by NaHCO₃. The m^7^G mutation rate (magenta) increases in the dual-treated sample compared to the other nucleotides (gray). Unlike 23S rRNA, 16S rRNA lacks D and OH^5^C modifications. D) The same experiment as in (C), but 16S rRNA was treated with KMnO₄ for 6 minutes instead of 3 minutes. E) Mutation rates of m^7^G, D, and OH^5^C modifications in 23S rRNA under various treatment conditions: 3 minutes with KMnO₄ (magenta), 6 minutes with KMnO₄ (blue), 3 minutes with KMnO₄ followed by NaHCO₃ (brown), and 6 minutes with KMnO₄ followed by NaHCO₃ (pink). F) Mutation rates of m^7^G modification in 16S rRNA treated with KMnO₄ alone or KMnO₄ followed by NaHCO₃, as described in (E). For all panels, mutation rates are background-corrected using Equation 3, and errors in panels (E) and (F) are propagated using Equation 5. In all panels, “nt” refers to nucleotides and “min” to minutes.

For the rRNA samples subjected to KMnO_4_ and KMnO_4_ plus NaHCO₃, we observe a larger number of false-positive modifications than in other conditions interrogated here (Table S3). KMnO_4_ has been found to oxidize G, T, and C bases, and less frequently A bases ^66, 67^. This oxidation process has been shown to produce polymerase mistakes ^68–72^. The false-positive modifications that we observe, which are more extensive in G bases, are likely due to G oxidation by KMnO_4_ (Table S3) ^66, 67^. Future experiments employing a smaller concentration of KMnO_4_ or exposing the RNA to KMnO_4_ for a shorter period of time could decrease the number of false-positive G modifications. In its present form, comparing the NGS data of RNA subjected to KMnO_4_ plus NaHCO₃ to those of RNA treated with CMCT plus NaHCO₃ will validate m^7^G and OH^5^C modifications and will discriminate between D and Ψ modifications.

### The RlmA enzyme, which incorporates m^1^G 747; the RlmG enzyme, which incorporates, m^2^G 1837; and the RlmL enzyme, which incorporates m^2^G 2449, have performed their functions before the 27S, 35S, and 45S accumulate in cells

In the untreated 23S rRNA samples, modified nucleotides m^1^G 747, m^2^G 1837, and m^2^G 2449 show similar levels of mutation rates across the three different intermediates, 27S, 35S, 45S and 50S (Figure 4). Since the 27S particle is a very early-stage large-subunit intermediate, in the pathway where the 27S accumulates, the RlmA, RlmG, and RlmL methyltransferases act during the very early stages of large subunit assembly. Similarly, the 35S intermediate is an early-stage large-subunit intermediate; thus, in the large subunit ribosome assembly pathway, where the 35S intermediate is populated, RlmA, RlmG, and RlmL could act during the very early or early stages of large subunit ribosome assembly. Lastly, the 45S is a late-stage large-subunit intermediate; therefore, in the large subunit ribosome assembly pathway where the 45S is populated, RlmA, RlmG, and RlmL could act during any of the very early, early, or intermediate stages of large subunit ribosome assembly.

**Figure 4.**
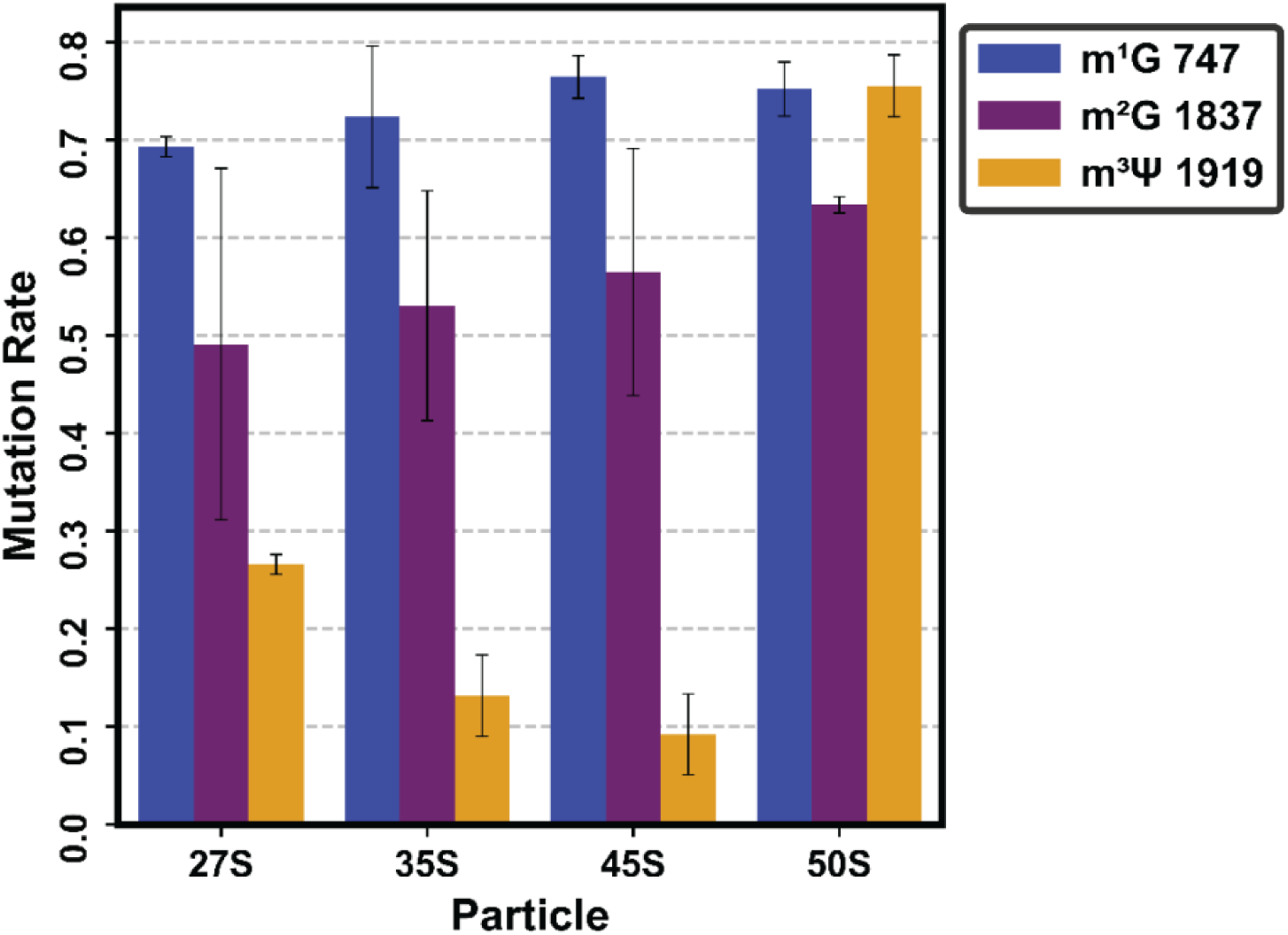
The m^1^G and m^2^G modifications are incorporated at similar levels in the 27S, 35S, and 45S intermediates, as well as in the 50S large subunit. The m^3^Ψ modification is incorporated to a lesser extent in the 27S, 35S, and 45S than in the 50S large subunit. The mutation rates shown here were determined from 23S rRNA samples that were chemically untreated. The values shown are the averages determined from two biological replicates (Equation 1); the errors represent the standard deviations from the averages (Equation 2). The mutation rate for m^1^G 747 is shown as blue bars, m^2^G 1837 as magenta, m^2^G 2449 as purple, and m^3^Ψ 1919 as brown. The m^3^Ψ modification is incorporated to a lesser extent in the 35S, an early-stage large-subunit assembly intermediate, and the 45S, a late-stage large-subunit assembly intermediate, when compared to the very-early-stage large-subunit ribosome assembly intermediate, 27S.

Previous studies have found that in *E. coli* cells exposed to erythromycin and chloramphenicol, in cells lacking the DEAD-box RNA helicase SrmB, and in wild-type cells, the RlmA and RlmG methyltransferases function during the early stages of large subunit ribosome assembly ^52–54^. Furthermore, in cells exposed to erythromycin or chloramphenicol, the RlmL enzyme was found to act during the early stages of large subunit ribosome assembly ^52^. However, no distinctions were made in the above studies between very-early and early stages of large-subunit ribosome assembly ^52–54^. Thus, in our cells expressing the helicase-inactive R331A DbpA construct, in the pathways where the 27S and 35S intermediates accumulate, the RlmA, RlmG, and RlmL enzymes act during the early stages of ribosome assembly, similar to previously investigated cellular or stress conditions ^52–54^.

The RlmL methyltransferase makes up one domain of the RlmKL protein; the other domain consists of the RlmK methyltransferase, which incorporates m^7^G 2073 modifications ^73^. The specific stage of ribosome assembly at which RlmK performs its function in cells expressing the R331A construct DbpA will be discussed in the next section.

Interestingly, we observe that while the m^3^Ψ 1919 modification is incorporated to a lesser extent in the 27S, 35S, and 45S intermediates compared to the wild-type 50S, this modification is incorporated to a greater extent in the very-early large subunit stage intermediate 27S than in the later-stage intermediates 35S and 45S (Figure 4). The m^3^Ψ 1919 modification is facilitated by the combined action of two enzymes (Table S1). First, the pseudouridine synthase RluD isomerizes U 1919 to Ψ ^74^. Subsequently, the RlmH methyltransferase adds a methyl group at the N3 position of Ψ 1919 ^25, 75^. Studies from various groups reveal that RluD and RlmH function both *in vivo* and *in vitro* during the late stages of ribosome assembly ^5, 25, 42, 43, 52–54, 76^. The Cryogenic Electron Microscopy analysis of a late-stage 50S ribosomal intermediate reveals that the RluD enzyme interacts with another maturation factor, YjgA, which is also involved in the late stages of ribosome assembly ^5, 77^. It is possible that certain unique RNA structures or early-acting maturation factors present in the pathway where the 27S intermediate accumulates, partially stimulate the isomerase function of RluD and the subsequent methyltransferase function of RlmH at position 1919 of 23S rRNA, compared to the 35S and 45S intermediates. The structure of the 27S intermediate remains unknown. Future structural studies could shed light on the RNA structures and maturation factor compositions that render the 27S intermediate more desirable for the RluD and RlmH enzymes to act upon than the 35S and 45S intermediates.

### The RlmK enzyme, which incorporates m^7^G 2073, and the RdsA enzyme, which incorporates D 2453, have carried out their respective functions in the 27S, 35S, and 45S intermediates

The mutation rates of m^7^G 2073 and D 2453 nucleotides in the 23S rRNA samples isolated from 27S, 35S, and 45S intermediates, as well as the mature wild-type 50S, are similar for samples treated with NaHCO_3_ or CMCT plus NaHCO_3_ (Figure 5A, and Figure 5B). Thus, the RlmK enzyme, which incorporates the m^7^G 2073 modification, and the RdsA enzyme, which incorporates the D 2453 modification, have carried out their functions in the 27S, 35S and 45S intermediates. Since the 27S particle is a very-early large-subunit intermediate, in the pathway in which it accumulates, the RlmK and RdsA enzymes act during the very-early stages of large-subunit assembly. Similarly, since the 35S intermediate is an early-stage large-subunit intermediate in the large-subunit ribosome assembly pathway, where it is populated, the RlmK and RdsA enzymes act during the very early or early stages of large-subunit ribosome assembly. Lastly, the 45S is a late-stage large-subunit intermediate; hence, in the large-subunit ribosome assembly pathway where it is populated, the RlmK and RdsA enzymes act during the very-early, early, or intermediate stages of large-subunit ribosome assembly.

**Figure 5.**
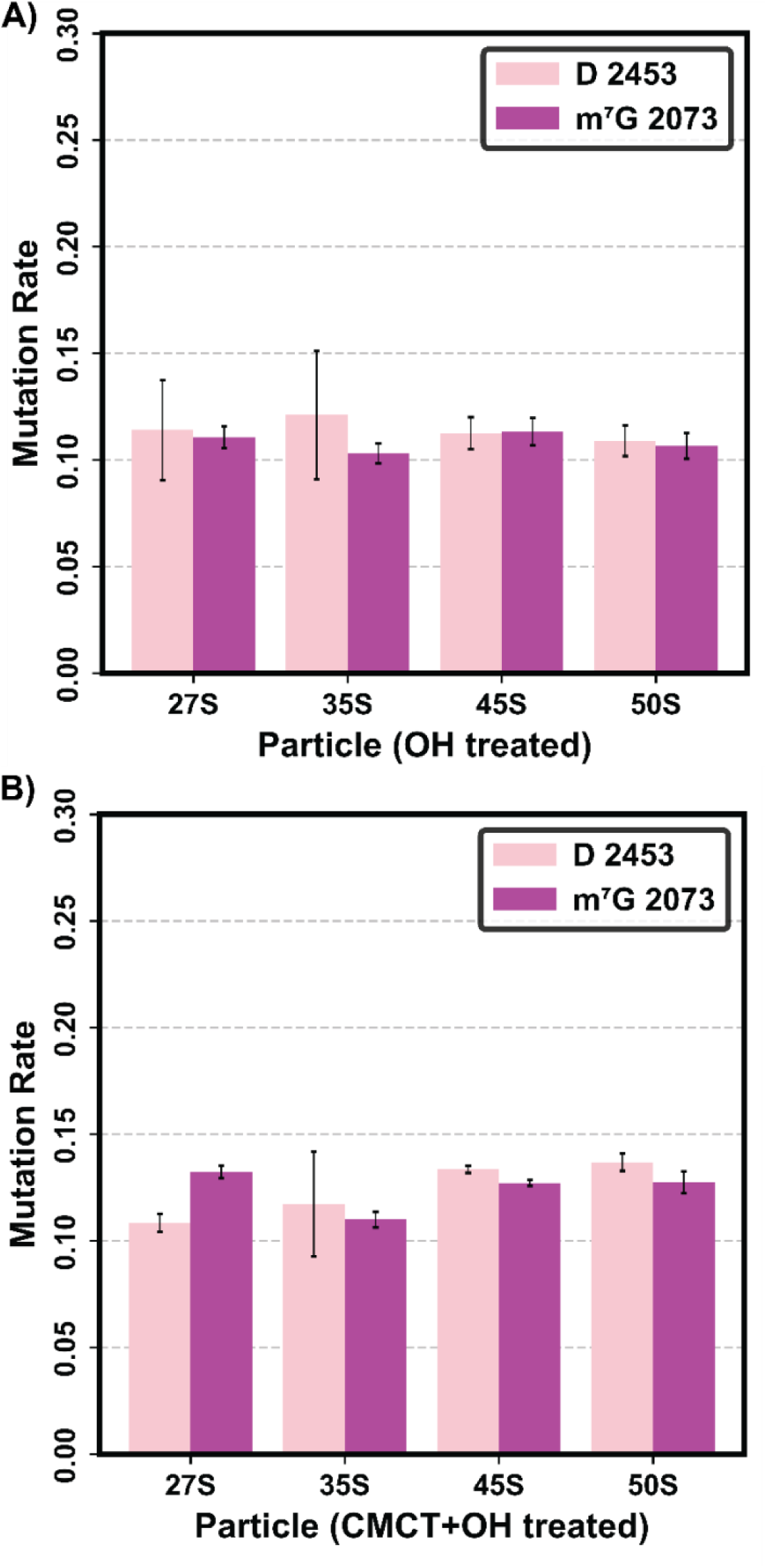
m^7^G and D are incorporated at similar levels in the 27S, 35S, and 45S large-subunit intermediates, as well as in the wild-type 50S. A) The mutation rates of D (pink) and m^7^G (purple) in the 27S, 35S, and 45S intermediates, as well as in the 50S large subunit, were determined by NaHCO3 treatment. Background corrections were performed using Equation 3, and errors were calculated using Equation 5. B) The mutation rates of D and m^7^G in the 27S, 35S, and 45S intermediates, as well as the 50S large subunit, were determined by CMCT and NaHCO_3_ treatments. The mutation rate values represent the averages from two biological replicates (Equation 7), and the errors indicate deviations from these averages (Equation 8). Subjecting the 23S rRNA samples to CMCT treatment, in addition to NaHCO_3_, has no effect on the D and m^7^G mutation rates in the intermediates or the 50S large subunit.

The RdsA enzyme was only recently identified, and the stage at which it acts during *E. coli* large-subunit ribosome assembly remained undetermined until this study ^52–54, 78^. A recent study found that the D modification is present in a late large-subunit ribosomal particle with a sedimentation coefficient of 45S, which accumulates in *Bacillus subtilis* (*B. subtilis*), at the same level as in the properly matured 50S ^79^. Thus, in *B. subtilis*, the D modification is incorporated during the early or intermediate stages of large-subunit ribosome assembly, consistent with the results presented in this study for *E. coli* large-subunit ribosome assembly (Figure 5) ^79^.

The RlmK enzyme was found to act during the early to intermediate stages of large-subunit ribosome assembly in *E. coli* cells exposed to erythromycin or chloramphenicol, and during the early stages of large-subunit ribosome assembly in wild-type cells or cells lacking the SrmB protein ^52–54^. Thus, in cells expressing the helicase-inactive R331A DbpA, in the pathways where the 27S and 35S intermediates are populated, the RlmK enzyme acts at similar time points during ribosome assembly as in wild-type cells or cells lacking the SrmB protein ^53, 54^. Moreover, it is possible that in cells expressing R331A DbpA, in the pathway where the 45S intermediate is populated, the RlmK enzyme could act during the early to intermediate stages of large-subunit ribosome assembly, similarly to cells exposed to erythromycin or chloramphenicol ^52^.

## CONCLUSIONS

In this work, we developed three Illumina NGS techniques to simultaneously detect multiple classes of RNA modifications: one without chemical treatment, one using CMCT plus alkaline conditions, and one using KMnO_4_ plus alkaline conditions. These techniques allowed us to detect the following rRNA modifications: m^1^G, m^2^G, m^3^Ψ, m^3^U, and m^6^ A (no chemical treatment; Figure 1); m^7^G, D, m^2^A, OH^5^C, and Ψ (CMCT plus alkaline; Figure 2); and m^7^G, D, and OH^5^C (KMnO_4_ plus alkaline; Figure 3). A comparative analysis of data from the CMCT plus alkaline treatment and the KMnO_4_ plus alkaline treatment could be used to validate the presence of m^7^G and OH^5^C modifications, while distinguishing between D and Ψ modifications in rRNA molecules with unknow modification compositions.

We applied these techniques to detect the presence of m^1^G, m^2^G, m^7^G, and D modifications in the 27S, 35S, and 45S intermediates that accumulate in cells expressing the R331A DbpA construct. Our results reveal that the m^1^G 747, m^2^G 1837, m^2^G 2449, m^7^G 2073, and D 2453 modifications are incorporated into these three intermediates at levels comparable to those in the mature 50S. Thus, the enzymes responsible for incorporating these modifications—RlmA, RlmG, RlmKL, and RdsA—complete their functions prior to the accumulation of the 27S, 35S, and 45S intermediates in cells (Figure 6).

**Figure 6.**
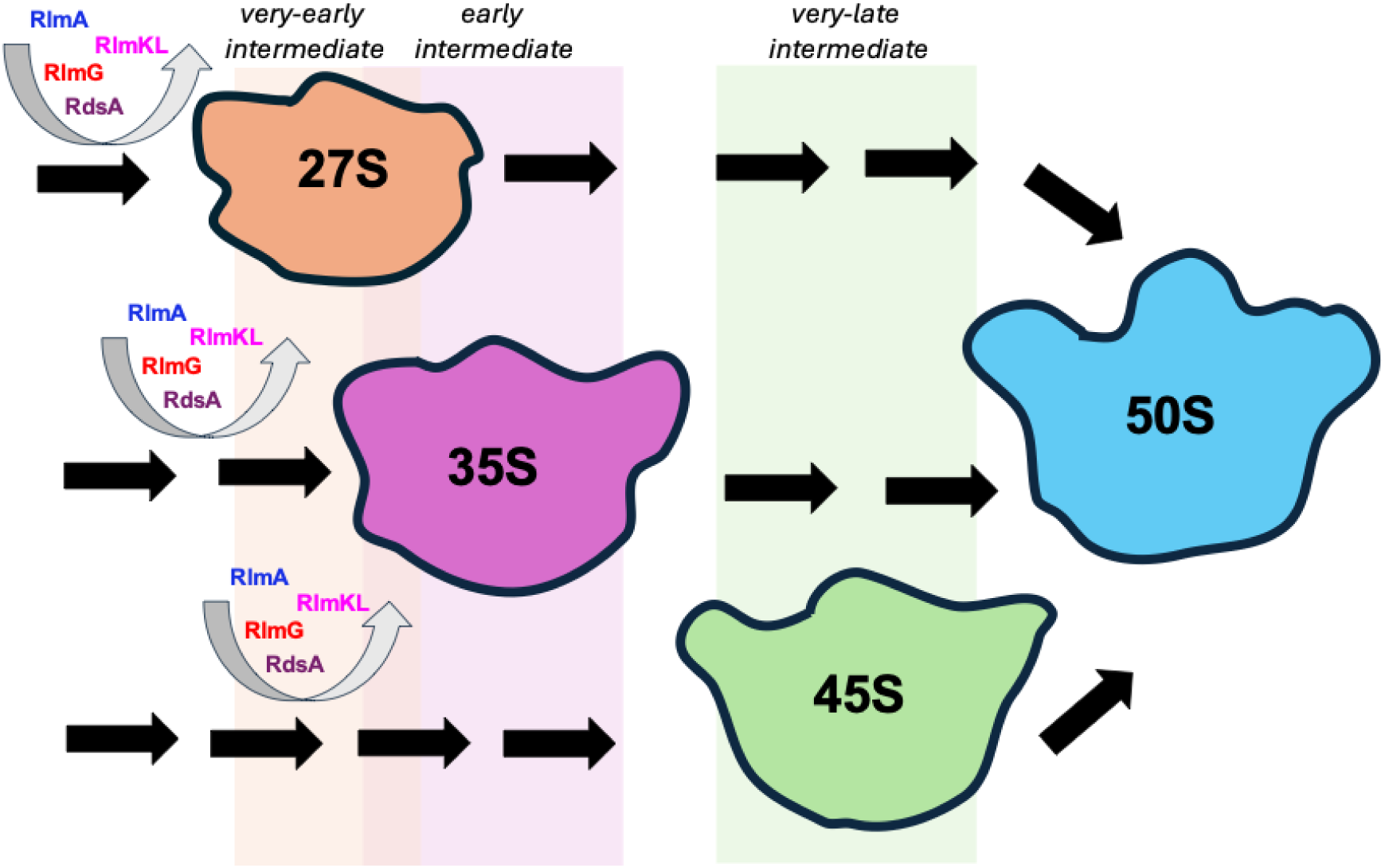
Summary of the timing of modification enzyme activity in three pathways of large ribosomal subunit assembly in cells expressing the R331A DbpA construct. The 27S is a very-early-stage large-subunit intermediate, as RdsA, RlmKL, RlmG, and RlmA perform their functions before the 27S accumulates in cells. In the pathway where the 27S intermediate is populated, RdsA, RlmKL, RlmG, and RlmA act at the very-early stages of large-subunit ribosome assembly (Table 1). The 35S is an early-stage large subunit intermediate, as RdsA, RlmKL, RlmG, and RlmA perform their functions before the 35S accumulates in cells. In the pathway where the 35S intermediate is populated, RdsA, RlmKL, RlmG, and RlmA act at the early stages of large subunit ribosome assembly (Table 1). The 45S is a late-stage large-subunit intermediate, as RdsA, RlmKL, RlmG, and RlmA perform their functions before the 45S accumulates in cells. In the pathway where the 45S intermediate is populated, RdsA, RlmKL, RlmG, and RlmA perform their enzymatic functions at the very-early, early, or intermediate stages of large subunit ribosome assembly (Table 1).

The 27S, 35S, and 45S intermediates belong to distinct pathways and stages of large-subunit ribosome assembly: very-early, early, and late (Figure 6). Our findings also reveal that the four enzymes—RlmA, RlmG, RlmKL, and RdsA—act in the pathway where the 27S intermediate accumulates during the very-early stages of large subunit ribosome assembly, in the pathway where the 35S intermediate accumulates during the early stages, and in the pathway where the 45S intermediate accumulates during the very-early, early, or intermediate stages of large-subunit ribosome assembly (Figure 6).

Finally, although the precise timing of RdsA-mediated reduction of U 2453 to D during *E. coli* large-subunit ribosome assembly was previously unknown, our study identifies the specific time points within the three pathways of large subunit ribosome assembly where the RdsA enzyme acts in cells expressing the R331A DbpA construct ^52–54^.

## DATA AVAILABILITY

The Illumina NGS raw files and ShapeMapper v1.2 output files were deposited on Gene Expression Omnibus (GEO). The GEO accession codes are: GSE196821; GSE232539; GSEXXXXXX

## ASSOCIATED CONTENT

### Supporting Information

List of positions of RNA modified nucleotides as well as the enzymes that insert them in 16S rRNA and 23S rRNA. Summary of NGS Illumina Techniques employed to detect RNA modifications. List of nucleotides erroneously marked as modified sites as per our 0.05 mutation rate threshold. The mismatch and deletion pattern of the reverse transcriptase at m^2^G modifications is sequence-context-independent and distinct from that of m^1^G. The mismatch and deletion profile of m^7^G is sequence-context-dependent. The error signatures of reverse transcriptase differ depending on the sequence context of Ψ modifications. D and m^7^G mutation rates are unaffected by KMnO4, whereas they increase in the presence of NaHCO_3_.

## Accession Code

*E.coli* DbpA Uniprot entry: P21693 E.coli SrmB Uniprot entery: P21507 *E.coli* RluD Uniprot entry: P33643 *E.coli* RlmH Uniprot entry: P0A818 *E.coli* RlmA Uniprot entry: P36999 *E.coli* RlmG Uniprot entry: P42596 *E.coli* RlmKL Uniprot entry: Q8XDB2 E.coli RdsA Uniprot entry: P37631

## FUNDING

This work was supported in part by the National Institute of General Medical Sciences grant R01-GM131062, the University of Texas System Rising STARs Program, and the start-up from the Chemistry and Biochemistry Department at the University of Texas at El Paso (to E.K.).

## ACKNOWLEDGMENTS

We are grateful to Riley C. Gentry, David Mohr, the Johns Hopkins University Genetic Core Facility and the University of Florida Interdisciplinary Center for Biotechnology Research for the help with library preparation and NGS data collection.

## FOR TABLE OF CONTENTS USE ONLY

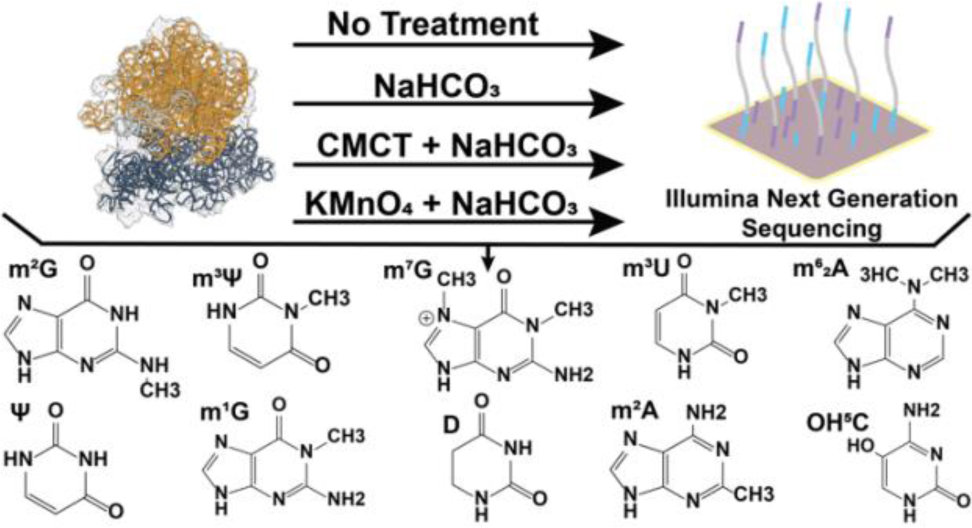

## Notes

### Competing Interest Statement

The authors have declared no competing interest.

## REFERENCES

[1] Toward Sequencing and Mapping of RNA Modifications Committee (Washington District of Columbia) (2024) Charting a future for sequencing rna and its modifications: a new era for biology and medicine, National Academies Press, Washington.

[2] Ero, R., Leppik, M., Reier, K., Liiv, A., and Remme, J. (2024) Ribosomal RNA modification enzymes stimulate large ribosome subunit assembly in E. coli, Nucleic Acids Res 52, 6614–6628.

[3] Boo, S. H., and Kim, Y. K. (2020) The emerging role of RNA modifications in the regulation of mRNA stability, Exp Mol Med 52, 400–408.

[4] Gilbert, W. V., and Nachtergaele, S. (2023) mRNA Regulation by RNA Modifications, Annu Rev Biochem 92, 175–198.

[5] Nikolay, R., Hilal, T., Schmidt, S., Qin, B., Schwefel, D., Vieira-Vieira, C. H., Mielke, T., Burger, J., Loerke, J., Amikura, K., Flugel, T., Ueda, T., Selbach, M., Deuerling, E., and Spahn, C. M. T. (2021) Snapshots of native pre-50S ribosomes reveal a biogenesis factor network and evolutionary specialization, Mol Cell, 1200–1215.

[6] Arai, T., Ishiguro, K., Kimura, S., Sakaguchi, Y., Suzuki, T., and Suzuki, T. (2015) Single methylation of 23S rRNA triggers late steps of 50S ribosomal subunit assembly, Proc Natl Acad Sci U S A 112, E4707–4716.

[7] Kimura, S., Ikeuchi, Y., Kitahara, K., Sakaguchi, Y., Suzuki, T., and Suzuki, T. (2012) Base methylations in the double-stranded RNA by a fused methyltransferase bearing unwinding activity, Nucleic Acids Res 40, 4071–4085.

[8] Fleming, A. M., Bommisetti, P., Xiao, S., Bandarian, V., and Burrows, C. J. (2023) Direct Nanopore Sequencing for the 17 RNA Modification Types in 36 Locations in the E. coli Ribosome Enables Monitoring of Stress-Dependent Changes, ACS Chem Biol.

[9] Fasnacht, M., Gallo, S., Sharma, P., Himmelstoss, M., Limbach, P. A., Willi, J., and Polacek, N. (2022) Dynamic 23S rRNA modification ho5C2501 benefits Escherichia coli under oxidative stress, Nucleic Acids Res 50, 473–489.

[10] Watkins, C. P., Zhang, W., Wylder, A. C., Katanski, C. D., and Pan, T. (2022) A multiplex platform for small RNA sequencing elucidates multifaceted tRNA stress response and translational regulation, Nat Commun 13, 2491.

[11] Babosan, A., Fruchard, L., Krin, E., Carvalho, A., Mazel, D., and Baharoglu, Z. (2022) Nonessential tRNA and rRNA modifications impact the bacterial response to sub-MIC antibiotic stress, microLife 3, 1–18.

[12] Vila-Sanjurjo, A., Squires, C. L., and Dahlberg, A. E. (1999) Isolation of kasugamycin resistant mutants in the 16 S ribosomal RNA of Escherichia coli, J Mol Biol 293, 1–8.

[13] Okamoto, S., Tamaru, A., Nakajima, C., Nishimura, K., Tanaka, Y., Tokuyama, S., Suzuki, Y., and Ochi, K. (2007) Loss of a conserved 7-methylguanosine modification in 16S rRNA confers low-level streptomycin resistance in bacteria, Mol Microbiol 63, 1096–1106.

[14] Delgado-Tejedor, A., Medina, R., Begik, O., Cozzuto, L., Lopez, J., Blanco, S., Ponomarenko, J., and Novoa, E. M. (2024) Native RNA nanopore sequencing reveals antibiotic-induced loss of rRNA modifications in the A- and P-sites, Nat Commun 15, 10054.

[15] Boccaletto, P., and Baginski, B. (2021) MODOMICS: An Operational Guide to the Use of the RNA Modification Pathways Database, Methods Mol Biol 2284, 481–505.

[16] Motorin, Y., and Helm, M. (2024) General Principles and Limitations for Detection of RNA Modifications by Sequencing, Acc Chem Res 57, 275–288.

[17] Khoddami, V., Yerra, A., Mosbruger, T. L., Fleming, A. M., Burrows, C. J., and Cairns, B. R. (2019) Transcriptome-wide profiling of multiple RNA modifications simultaneously at single-base resolution, Proc Natl Acad Sci U S A 116, 6784–6789.

[18] Araujo Tavares, R. C., Mahadeshwar, G., Wan, H., and Pyle, A. M. (2023) MRT-ModSeq - Rapid Detection of RNA Modifications with MarathonRT, J Mol Biol 435, 168299.

[19] Noeske, J., Wasserman, M. R., Terry, D. S., Altman, R. B., Blanchard, S. C., and Cate, J. H. (2015) High-resolution structure of the Escherichia coli ribosome, Nat Struct Mol Biol 22, 336–341.

[20] Shajani, Z., Sykes, M. T., and Williamson, J. R. (2011) Assembly of bacterial ribosomes, Annu Rev Biochem 80, 501–526.

[21] Sergiev, P. V., Lesnyak, D. V., Bogdanov, A. A., and Dontsova, O. A. (2006) Identification of Escherichia coli m2G methyltransferases: II. The ygjO Gene Encodes a Methyltransferase Specific for G1835 of the 23 S rRNA, Journal of Molecular Biology 364, 26–31.

[22] Sergiev, P. V., Serebryakova, M. V., Bogdanov, A. A., and Dontsova, O. A. (2008) The ybiN Gene of Escherichia coli Encodes Adenine-N6 Methyltransferase Specific for Modification of A1618 of 23 S Ribosomal RNA, a Methylated Residue Located Close to the Ribosomal Exit Tunnel, Journal of Molecular Biology 375, 291–300.

[23] Ofengand, J., and Del Campo, M. (2004) Modified Nucleosides of Escherichia coli Ribosomal RNA, EcoSal Plus 1.

[24] Havelund, J. F., Giessing, A. M., Hansen, T., Rasmussen, A., Scott, L. G., and Kirpekar, F. (2011) Identification of 5-hydroxycytidine at position 2501 concludes characterization of modified nucleotides in E. coli 23S rRNA, J Mol Biol 411, 529–536.

[25] Ero, R., Peil, L., Liiv, A., and Remme, J. (2008) Identification of pseudouridine methyltransferase in Escherichia coli, RNA 14, 2223–2233.

[26] Purta, E., O’Connor, M., Bujnicki, J. M., and Douthwaite, S. (2008) YccW is the m5C Methyltransferase Specific for 23S rRNA Nucleotide 1962, Journal of Molecular Biology 383, 641–651.

[27] Wang, K.-T., Desmolaize, B., Nan, J., Zhang, X.-W., Li, L.-F., Douthwaite, S., and Su, X.-D. (2012) Structure of the bifunctional methyltransferase YcbY (RlmKL) that adds the m 7 G2069 and m 2 G2445 modifications in Escherichia coli 23S rRNA, Nucleic Acids Research 40, 5138–5148.

28. Toubdji, S., Thullier, Q., Kilz, L.-M., Marchand, V., Yuan, Y., Sudol, C., Goyenvalle, C., Jean-Jean, O., Rose, S., Douthwaite, S., Hardy, L., Baharoglu, Z., De Crécy-Lagard, V., Helm, M., Motorin, Y., Hamdane, D., and Brégeon, D. (2024) Exploring a unique class of flavoenzymes: Identification and biochemical characterization of ribosomal RNA dihydrouridine synthase, Proceedings of the National Academy of Sciences 121.

29. Purta, E., O’Connor, M., Bujnicki, J. M., and Douthwaite, S. (2009) YgdE is the 2′-O-ribose methyltransferase RlmM specific for nucleotide C2498 in bacterial 23S rRNA, Molecular Microbiology 72, 1147–1158.

[30] Kimura, S., Sakai, Y., Ishiguro, K., and Suzuki, T. (2017) Biogenesis and iron-dependency of ribosomal RNA hydroxylation, Nucleic Acids Research 45, 12974–12986.

[31] Toh, S.-M., Xiong, L., Bae, T., and Mankin, A. S. (2008) The methyltransferase YfgB/RlmN is responsible for modification of adenosine 2503 in 23S rRNA, RNA 14, 98–106.

[32] Okamoto, S., Tamaru, A., Nakajima, C., Nishimura, K., Tanaka, Y., Tokuyama, S., Suzuki, Y., and Ochi, K. (2007) Loss of a conserved 7-methylguanosine modification in 16S rRNA confers low-level streptomycin resistance in bacteria, Molecular Microbiology 63, 1096–1106.

[33] Lesnyak, D. V., Osipiuk, J., Skarina, T., Sergiev, P. V., Bogdanov, A. A., Edwards, A., Savchenko, A., Joachimiak, A., and Dontsova, O. A. (2007) Methyltransferase That Modifies Guanine 966 of the 16 S rRNA, Journal of Biological Chemistry 282, 5880–5887.

[34] Kimura, S., and Suzuki, T. (2010) Fine-tuning of the ribosomal decoding center by conserved methyl-modifications in the Escherichia coli 16S rRNA, Nucleic Acids Research 38, 1341–1352.

[35] Andersen, N. M., and Douthwaite, S. (2006) YebU is a m5C Methyltransferase Specific for 16 S rRNA Nucleotide 1407, Journal of Molecular Biology 359, 777–786.

[36] Basturea, G. N., Rudd, K. E., and Deutscher, M. P. (2006) Identification and characterization of RsmE, the founding member of a new RNA base methyltransferase family, RNA 12, 426–434.

[37] Basturea, G. N., Dague, D. R., Deutscher, M. P., and Rudd, K. E. (2012) YhiQ Is RsmJ, the Methyltransferase Responsible for Methylation of G1516 in 16S rRNA of E. coli, Journal of Molecular Biology 415, 16–21.

[38] Gc, K., Gyawali, P., Balci, H., and Abeysirigunawardena, S. (2020) Ribosomal RNA Methyltransferase RsmC Moonlights as an RNA Chaperone, Chembiochem 21, 1885–1892.

[39] Marchand, V., Bourguignon-Igel, V., Helm, M., and Motorin, Y. (2021) Mapping of 7-methylguanosine (m(7)G), 3-methylcytidine (m(3)C), dihydrouridine (D) and 5-hydroxycytidine (ho(5)C) RNA modifications by AlkAniline-Seq, Methods Enzymol 658, 25–47.

[40] Gentry, R. C., Childs, J. J., Gevorkyan, J., Gerasimova, Y. V., and Koculi, E. (2016) Time course of large ribosomal subunit assembly in E. coli cells overexpressing a helicase inactive DbpA protein, RNA 22, 1055–1064.

[41] Elles, L. M. S., Sykes, M. T., Williamson, J. R., and Uhlenbeck, O. C. (2009) A dominant negative mutant of the E. coli RNA helicase DbpA blocks assembly of the 50S ribosomal subunit, Nucleic Acids Res 37, 6503–6514.

[42] Koculi, E., and Cho, S. S. (2022) RNA Post-Transcriptional Modifications in Two Large Subunit Intermediates Populated in E. coli Cells Expressing Helicase Inactive R331A DbpA, Biochemistry 61, 833–842.

[43] Narayan, G., Gracia Mazuca, L. A., Cho, S. S., Mohl, J. E., and Koculi, E. (2023) RNA Post-transcriptional Modifications of an Early-Stage Large-Subunit Ribosomal Intermediate, Biochemistry 62, 2908–2915.

[44] Fuller-Pace, F. V., Nicol, S. M., Reid, A. D., and Lane, D. P. (1993) DbpA: a DEAD box protein specifically activated by 23s rRNA, Embo J 12, 3619–3626.

[45] Nicol, S. M., and Fuller-Pace, F. V. (1995) The “DEAD box” protein DbpA interacts specifically with the peptidyltransferase center in 23S rRNA, Proc Natl Acad Sci U S A 92, 11681–11685.

[46] Tsu, C. A., and Uhlenbeck, O. C. (1998) Kinetic analysis of the RNA-dependent adenosinetriphosphatase activity of DbpA, an Escherichia coli DEAD protein specific for 23S ribosomal RNA, Biochemistry 37, 16989–16996.

[47] Linder, P., and Jankowsky, E. (2011) From unwinding to clamping - the DEAD box RNA helicase family, Nat Rev Mol Cell Biol 12, 505–516.

[48] Childs, J. J., Gentry, R. C., Moore, A. F., and Koculi, E. (2016) The DbpA catalytic core unwinds double-helix substrates by directly loading on them, RNA 22, 408–415.

[49] Moore, A. F., Gentry, R. C., and Koculi, E. (2017) DbpA is a region-specific RNA helicase, Biopolymers 107.

50. Lopez de Victoria, A., Moore, A. F. T., Gittis, A. G., and Koculi, E. (2017) Kinetics and Thermodynamics of DbpA Protein’s C-Terminal Domain Interaction with RNA, ACS Omega 2, 8033–8038.

[51] Bohnsack, K. E., Yi, S., Venus, S., Jankowsky, E., and Bohnsack, M. T. (2023) Cellular functions of eukaryotic RNA helicases and their links to human diseases, Nat Rev Mol Cell Biol.

[52] Siibak, T., and Remme, J. (2010) Subribosomal particle analysis reveals the stages of bacterial ribosome assembly at which rRNA nucleotides are modified, RNA 16, 2023–2032.

[53] Popova, A. M., and Williamson, J. R. (2014) Quantitative analysis of rRNA modifications using stable isotope labeling and mass spectrometry, J Am Chem Soc 136, 2058–2069.

[54] Rabuck-Gibbons, J. N., Popova, A. M., Greene, E. M., Cervantes, C. F., Lyumkis, D., and Williamson, J. R. (2020) SrmB Rescues Trapped Ribosome Assembly Intermediates, J Mol Biol 432, 978–990.

[55] Siegfried, N. A., Busan, S., Rice, G. M., Nelson, J. A., and Weeks, K. M. (2014) RNA motif discovery by SHAPE and mutational profiling (SHAPE-MaP), Nat Methods 11, 959–965.

[56] Langmead, B., and Salzberg, S. L. (2012) Fast gapped-read alignment with Bowtie 2, Nat Methods 9, 357–359.

[57] Busan, S., and Weeks, K. M. (2018) Accurate detection of chemical modifications in RNA by mutational profiling (MaP) with ShapeMapper 2, RNA 24, 143–148.

[58] Hauenschild, R., Tserovski, L., Schmid, K., Thuring, K., Winz, M. L., Sharma, S., Entian, K. D., Wacheul, L., Lafontaine, D. L., Anderson, J., Alfonzo, J., Hildebrandt, A., Jaschke, A., Motorin, Y., and Helm, M. (2015) The reverse transcription signature of N-1-methyladenosine in RNA-Seq is sequence dependent, Nucleic Acids Res 43, 9950–9964.

[59] Zhou, K. I., Clark, W. C., Pan, D. W., Eckwahl, M. J., Dai, Q., and Pan, T. (2018) Pseudouridines have context-dependent mutation and stop rates in high-throughput sequencing, RNA Biol 15, 892–900.

60. Liu-Wei, W., van der Toorn, W., Bohn, P., Holzer, M., Smyth, R. P., and von Kleist, M. (2024) Sequencing accuracy and systematic errors of nanopore direct RNA sequencing, BMC Genomics 25, 528.

[61] Batt, D. R., Martin, Keith. J., Ploeser, M. James, Murray, James (1954) Chemistry of the Dihydropyrimidines. Ultraviolet Spectra and Alkaline Decomposition, Journal of the American Chemical Society 76, 3663–3665.

[62] Xing, F., Hiley, S. L., Hughes, T. R., and Phizicky, E. M. (2004) The specificities of four yeast dihydrouridine synthases for cytoplasmic tRNAs, J Biol Chem 279, 17850–17860.

[63] Lawley, P. D., and Shah, S. A. (1972) Methylation of ribonucleic acid by the carcinogens dimethyl sulphate, N-methyl-N-nitrosourea and N-methyl-N’-nitro-N-nitrosoguanidine. Comparisons of chemical analyses at the nucleoside and base levels, Biochem J 128, 117–132.

[64] Wintermeyer, W., and Zachau, H. G. (1970) A specific chemical chain scission of tRNA at 7-methylguanosine, FEBS Lett 11, 160–164.

[65] Zhang, L. S., Ju, C. W., Liu, C., Wei, J., Dai, Q., Chen, L., Ye, C., and He, C. (2022) m(7)G-quant-seq: Quantitative Detection of RNA Internal N(7)-Methylguanosine, ACS Chem Biol 17, 3306–3312.

[66] Bui, C. T., Rees, K., and Cotton, R. G. (2003) Permanganate oxidation reactions of DNA: perspective in biological studies, Nucleosides Nucleotides Nucleic Acids 22, 1835–1855.

[67] Bui, C. T., and Cotton, R. G. (2002) Comparative study of permanganate oxidation reactions of nucleotide bases by spectroscopy, Bioorg Chem 30, 133–137.

[68] Sarasin, A., Bounacer, A., Lepage, F., Schlumberger, M., and Suarez, H. G. (1999) Mechanisms of mutagenesis in mammalian cells. Application to human thyroid tumours, C R Acad Sci III 322, 143–149.

[69] Kamiya, H., Miura, H., Murata-Kamiya, N., Ishikawa, H., Sakaguchi, T., Inoue, H., Sasaki, T., Masutani, C., Hanaoka, F., Nishimura, S., and, et al. (1995) 8-Hydroxyadenine (7,8-dihydro-8-oxoadenine) induces misincorporation in in vitro DNA synthesis and mutations in NIH 3T3 cells, Nucleic Acids Res 23, 2893–2899.

[70] Suzuki, T., and Kamiya, H. (2017) Mutations induced by 8-hydroxyguanine (8-oxo-7,8-dihydroguanine), a representative oxidized base, in mammalian cells, Genes Environ 39, 2.

[71] Kreutzer, D. A., and Essigmann, J. M. (1998) Oxidized, deaminated cytosines are a source of C --> T transitions in vivo, Proc Natl Acad Sci U S A 95, 3578–3582.

[72] Hori, M., Yonekura, S., Nohmi, T., Gruz, P., Sugiyama, H., Yonei, S., and Zhang-Akiyama, Q. M. (2010) Error-Prone Translesion DNA Synthesis by Escherichia coli DNA Polymerase IV (DinB) on Templates Containing 1,2-dihydro-2-oxoadenine, J Nucleic Acids 2010, 807579.

[73] Wang, K. T., Desmolaize, B., Nan, J., Zhang, X. W., Li, L. F., Douthwaite, S., and Su, X. D. (2012) Structure of the bifunctional methyltransferase YcbY (RlmKL) that adds the m7G2069 and m2G2445 modifications in Escherichia coli 23S rRNA, Nucleic Acids Res 40, 5138–5148.

[74] Huang, L., Ku, J., Pookanjanatavip, M., Gu, X., Wang, D., Greene, P. J., and Santi, D. V. (1998) Identification of two Escherichia coli pseudouridine synthases that show multisite specificity for 23S RNA, Biochemistry 37, 15951–15957.

[75] Ero, R., Leppik, M., Liiv, A., and Remme, J. (2010) Specificity and kinetics of 23S rRNA modification enzymes RlmH and RluD, RNA 16, 2075–2084.

[76] Leppik, M., Peil, L., Kipper, K., Liiv, A., and Remme, J. (2007) Substrate specificity of the pseudouridine synthase RluD in Escherichia coli, FEBS J 274, 5759–5766.

[77] Du, M., Deng, C., Yu, T., Zhou, Q., and Zeng, F. (2024) YjgA plays dual roles in enhancing PTC maturation, Nucleic Acids Res 52, 7947–7960.

78. Toubdji, S., Thullier, Q., Kilz, L. M., Marchand, V., Yuan, Y., Sudol, C., Goyenvalle, C., Jean-Jean, O., Rose, S., Douthwaite, S., Hardy, L., Baharoglu, Z., de Crecy-Lagard, V., Helm, M., Motorin, Y., Hamdane, D., and Bregeon, D. (2024) Exploring a unique class of flavoenzymes: Identification and biochemical characterization of ribosomal RNA dihydrouridine synthase, Proc Natl Acad Sci U S A 121, e2401981121.

[79] Popova, A. M., Jain, N., Dong, X., Abdollah-Nia, F., Britton, R. A., and Williamson, J. R. (2024) Complete list of canonical post-transcriptional modifications in the Bacillus subtilis ribosome and their link to RbgA driven large subunit assembly, Nucleic Acids Res 52, 11203–11217.

