## Supporting Information for "Post-transcriptional Modifications of the Large Ribosome Subunit Assembly Intermediates in *E. coli* Expressing Helicase-Inactive DbpA Variant"

**Table S1**. List of positions of RNA modified nucleotides as well as the enzymes that insert them in 16S ribosomal RNA (rRNA) and 23S rRNA

| **16S rRNA nt Position[a]** | **Modification** | **Enzyme** |
|---|---|---|
| 516 | $\Psi$ | RsuA[1] |
| 527 | $m^7G$ | RsmG[2] |
| 966 | $m^2G$ | RsmD[3] |
| 967 | $m^5C$ | RsmB[1] |
| 1207 | $m^2G$ | RsmC[1] |
| 1402 | $m^4C_m$ | RsmH, RsmL[b, 4] |
| 1407 | $m^5C$ | RsmF[5] |
| 1498 | $m^3U$ | RsmE[6] |
| 1516 | $m^2G$ | RsmJ[7] |
| 1518 | $m^6_2A$ | RsmA[1] |
| 1519 | $m^6_2A$ | RsmA[1] |
| **23S rRNA nt Position[a]** | **Modification** | **Enzyme** |
| 747 | $m^1G$ | RlmA[1] |
| 748 | $\Psi$ | RluA[1] |
| 749 | $m^5U$ | RlmC[1] |
| 957 | $\Psi$ | RluC[1] |
| 1620 | $m^6A$ | RlmF[8] |
| 1837 | $m^2G$ | RlmG[9] |
| 1915 | $\Psi$ | RluD[1] |
| 1919 | $m^3\Psi$ | RluD, RlmH[c, 1, 10] |
| 1921 | $\Psi$ | RluD[1] |
| 1943 | $m^5U$ | RlmD[1] |
| 1966 | $m^5C$ | RlmI[11] |
| 2034 | $m^6A$ | RlmJ[12] |
| 2073 | $m^7G$ | RlmKL[13] |
| 2255 | $G_m$ | RlmB |
| 2449 | $m^2G$ | RlmKL[13] |
| 2453 | D | RdsA[14] |
| 2461 | $\Psi$ | RluE[1] |
| 2502 | $C_m$ | RlmM[15] |
| 2505 | $OH^5C$ | RlhA[16] |
| 2507 | $m^2A$ | RlmN[17] |
| 2508 | $\Psi$ | RluC[1] |
| 2556 | $U_m$ | RlmE[1] |
| 2584 | $\Psi$ | RluC[1] |
| 2608 | $\Psi$ | RluF[1] |
| 2609 | $\Psi$ | RluB[1] |

[a]Positions of rRNA modifications are those corresponding to *rrlB* and *rrsB* genes that encode 23S rRNA and 16S rRNAs, respectively.

[b]RsmH and RsmL are responsible for the methylations at the N4 and 2'-O positions of C 1402 in the 16S rRNA, respectively [4].

[c]The RluD pseudouridine synthase incorporates Ψ 1919 into 23S rRNA, while the RlmH enzyme methylates Ψ 1919 at position 3, thereby creating $m^3Ψ$ [1, 10].

**Table S2**. Summary of NGS Illumina Techniques employed to detect RNA modifications

| Modification | Detection Method | Treatment/Enzyme | References |
|---|---|---|---|
| $m^1G$ | Stops detection/stops removal. | Demethylase treatment. | Clark *et al.* [18], Dai *et al.* [19] |
| $m^1G$ | Stops, mismatch, deletions. | Reverse transcriptase reaction performed in the presence of $Mn^{2+}$. | Kristen *et al.*[20], Werner *et al.*[21] |
| $m^2G$ | Stops. | | Dai *et al.* [19] |
| $m^3U$ | Stops detection/stops removal combined with demethylase treatment. | Demethylase treatment. | Clark *et al.* [18] |
| $m^3U$ | Insertions, deletions, mismatches.<br><br>Machine learning is required for detection. | MarathonRT reverse transcriptase. | Araujo Tavares *et al.*[22] |
| $m^6{}_2A$ | Stops, mismatch, deletion. | No treatment | Werner *et al.*[21] |
| $m^4C_m$ | None | None | |
| $m^3\Psi$ | Mismatches, deletions. | No treatment | Koculi *et al.*[23], Narayan *et al.*[24] |
| D | Stops. | $NaBH_4$ followed by rhodamine labeling. | Finet *et al.* [25] |
| D | Stops. | $NaBH_4$. | Draycott *et al.* [26] |
| D | Stops. | $OH^-$ followed by aniline treatment. | Marchand *et al.* [27] |
| $m^7G$ | Stops | $OH^-$ followed by aniline treatment. | Marchand *et al.* [27, 28] |
| $m^7G$ | Mismatches, deletions. | $NaBH_4$. | Enroth *et al.* [29] |
| $m^7G$ | Mismatches. | $NaBH_4$ plus biotin labeling. | Zhang *et al.*[30] |
| $m^7G$ | Mismatches. | $KBH_4$ followed by acidic conditions. | Zhang *et al.*[31] |
| $m^7G$ | Insertions, deletions, mismatches.<br><br>Machine learning is required for detection. | MarathonRT reverse transcriptase. | Araujo Tavares *et al.*[22] |
| $OH^5C$ | Stops. | $OH^-$ followed by aniline treatment | Marchand *et al.* [27] |
| $OH^5C$ | Mismatches, deletions. | CMCT treatment. | Narayan *et al.*[24] |

**Table S3**. List of nucleotides erroneously marked as modified sites as per our 0.05 mutation rate threshold

| Particle | Position[a] | Nucleotide | Mutation Rate[b] R1 | Mutation Rate[b] R2 | Chemical Treatment[c] |
|---|---|---|---|---|---|
| 30S | 271 | C | 0.149 | 0.024 | Untreated |
| | 856 | C | 0.124 | 0.005 | |
| | 867 | G | 0.079 | 0.004 | |
| | 6 | G | 0.059 | - | $OH^-$ |
| | 9 | G | 0.056 | - | |
| | 1515 | G | 0.053 | - | |
| | 6 | G | 0.085 | 0.154 | CMCT + $OH^-$ |
| | 9 | G | 0.077 | 0.131 | |
| | 1515 | G | 0.048 | 0.055 | |
| | 35 | G | 0.065 | - | $KMnO_4$ (3 min) |
| | 319 | G | 0.084 | - | |
| | 324 | G | 0.060 | - | |
| | 6 | G | 0.066 | - | $KMnO_4$ (3 min) + $OH^-$ |
| | 319 | G | 0.075 | - | |
| | 324 | G | 0.053 | - | |
| | 266 | G | 0.053 | - | $KMnO_4$ (6 min) |
| | 319 | G | 0.099 | - | |
| | 324 | G | 0.081 | - | |
| | 6 | G | 0.118 | - | $KMnO_4$ (6 min) + $OH^-$ |
| | 9 | G | 0.060 | - | |
| | 319 | G | 0.090 | - | |
| | 324 | G | 0.076 | - | |
| 50S | 1384 | G | 0.095 | 0.112 | Untreated |
| | 1660 | C | 0.125 | 0.051 | |
| | 1508 | T | 0.120 | - | $OH^-$ |
| | 829 | T | 0.058 | 0.053 | CMCT + $OH^-$ |
| | 1508 | T | 0.129 | 0.129 | |
| | 220 | G | 0.060 | - | $KMnO_4$ (3 min) |
| | 676 | G | 0.060 | - | |
| | 1225 | G | 0.173 | - | |
| | 1227 | G | 0.059 | - | |
| | 1273 | G | 0.055 | - | |
| | 1477 | G | 0.051 | - | |
| | 1525 | T | 0.092 | - | |
| | 1635 | G | 0.055 | - | |

| | | | | |
|---|---|---|---|---|
| 2367 | G | 0.059 | - | |
| 2894 | G | 0.108 | - | |
| 220 | G | 0.053 | - | $KMnO_4$ (3 min) + $OH^-$ |
| 676 | G | 0.063 | - | |
| 1225 | G | 0.155 | - | |
| 1227 | G | 0.052 | - | |
| 1273 | G | 0.058 | - | |
| 1508 | T | 0.060 | - | |
| 1525 | T | 0.151 | - | |
| 1635 | G | 0.053 | - | |
| 2894 | G | 0.093 | - | |
| 220 | G | 0.091 | - | $KMnO_4$ (6 min) |
| 372 | G | 0.053 | - | |
| 463 | G | 0.051 | - | |
| 661 | G | 0.050 | - | |
| 676 | G | 0.089 | - | |
| 829 | T | 0.055 | - | |
| 1225 | G | 0.252 | - | |
| 1227 | G | 0.075 | - | |
| 1273 | G | 0.076 | - | |
| 1477 | G | 0.073 | - | |
| 1508 | T | 0.055 | - | |
| 1525 | T | 0.108 | - | |
| 1635 | G | 0.080 | - | |
| 2367 | G | 0.085 | - | |
| 2599 | G | 0.064 | - | |
| 2622 | G | 0.064 | - | |
| 2894 | G | 0.151 | - | |
| 220 | G | 0.076 | - | $KMnO_4$ (6 min) + $OH^-$ |
| 372 | G | 0.052 | - | |
| 676 | G | 0.090 | - | |
| 1225 | G | 0.227 | - | |
| 1227 | G | 0.073 | - | |
| 1273 | G | 0.090 | - | |
| 1302 | G | 0.051 | - | |
| 1477 | G | 0.071 | - | |
| 1508 | T | 0.054 | - | |
| 1525 | T | 0.226 | - | |
| 1635 | G | 0.074 | - | |
| 1720 | G | 0.051 | - | |

50S

| | | | |
|---|---|---|---|
| 2367 | G | 0.072 | - |
| 2599 | G | 0.057 | - |
| 2622 | G | 0.059 | - |
| 2894 | G | 0.150 | - |

[a] These nucleotide positions correspond to the 16S rRNA *rrsB* gene and the 23S rRNA *rrlB* gene. The 16S and 23S rRNA molecules were extracted from the mature 30S and 50S ribosomal subunits, respectively.

[b] The mutation rates presented here correspond to nucleotides that are known to be unmodified in 16S and 23S rRNA and exhibit mutation rates greater than 0.05 (Table S1). In this study, a mutation rate threshold of 0.05 was used to identify detectable RNA modifications.

Mutation rates for all chemically treated samples were background-corrected using the average mutation rate of the corresponding nucleotide in untreated samples, as described in the Materials and Methods section.

Mutation rates for untreated samples were not background-corrected. R1 and R2 represent biological replicates 1 and 2, respectively

[c] 16S rRNA and 23S rRNA, as detailed in the manuscript, were either left untreated or subjected to the following chemical treatments: $NaHCO_3$; CMCT followed by $NaHCO_3$; $KMnO_4$ for 3 minutes; $KMnO_4$ for 6 minutes; $KMnO_4$ for 3 minutes followed by $NaHCO_3$; or $KMnO_4$ for 6 minutes followed by $NaHCO_3$. In this table, the abbreviation 'min' refers to minutes.

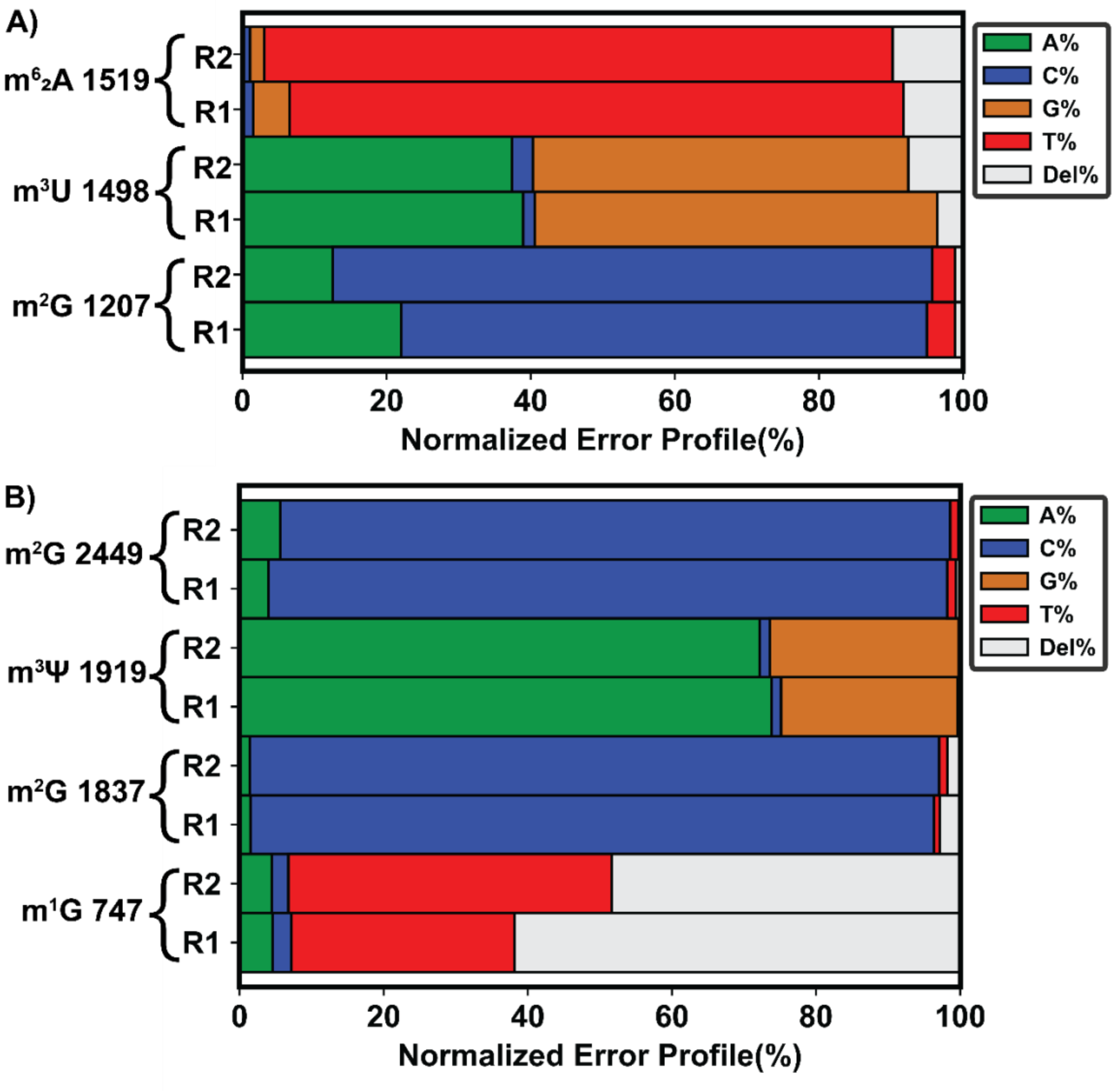

**Figure S1.** The mismatch and deletion pattern of the reverse transcriptase at m$^2$G modifications is sequence-context-independent and distinct from that of m$^1$G. Panels A (16S rRNA from the mature 30S) and B (23S rRNA from mature 50S) show m$^1$G-, m$^2$G-, m$^3$Ψ-, m$^3$U-, and m$^6_2$A- induced reverse transcriptase signatures of deletions and mismatches across the two biological replicates, R1 and R2. m$^2$G is the only modification detected in multiple sequences within 16S and 23S rRNA (Table 1 and Table 2). The reverse transcriptase signature profiles of the two detectable m$^2$G modifications are similar but differ from the m$^1$G modification (Panels A and B). For both panels the specific class of mismatches and deletions were depicted as the percentages of total reverse transcriptase errors, which was calculated by ShapeMapper v1.2. Legend: A green bar represents the percentage of the modified nucleotide miscalled as A, a blue bar represents the percentage miscalled as C, a brown bar represents the percentage miscalled as G, a red bar represents the percentage miscalled as T, and a gray bar represents the percentage identified as a deletion.

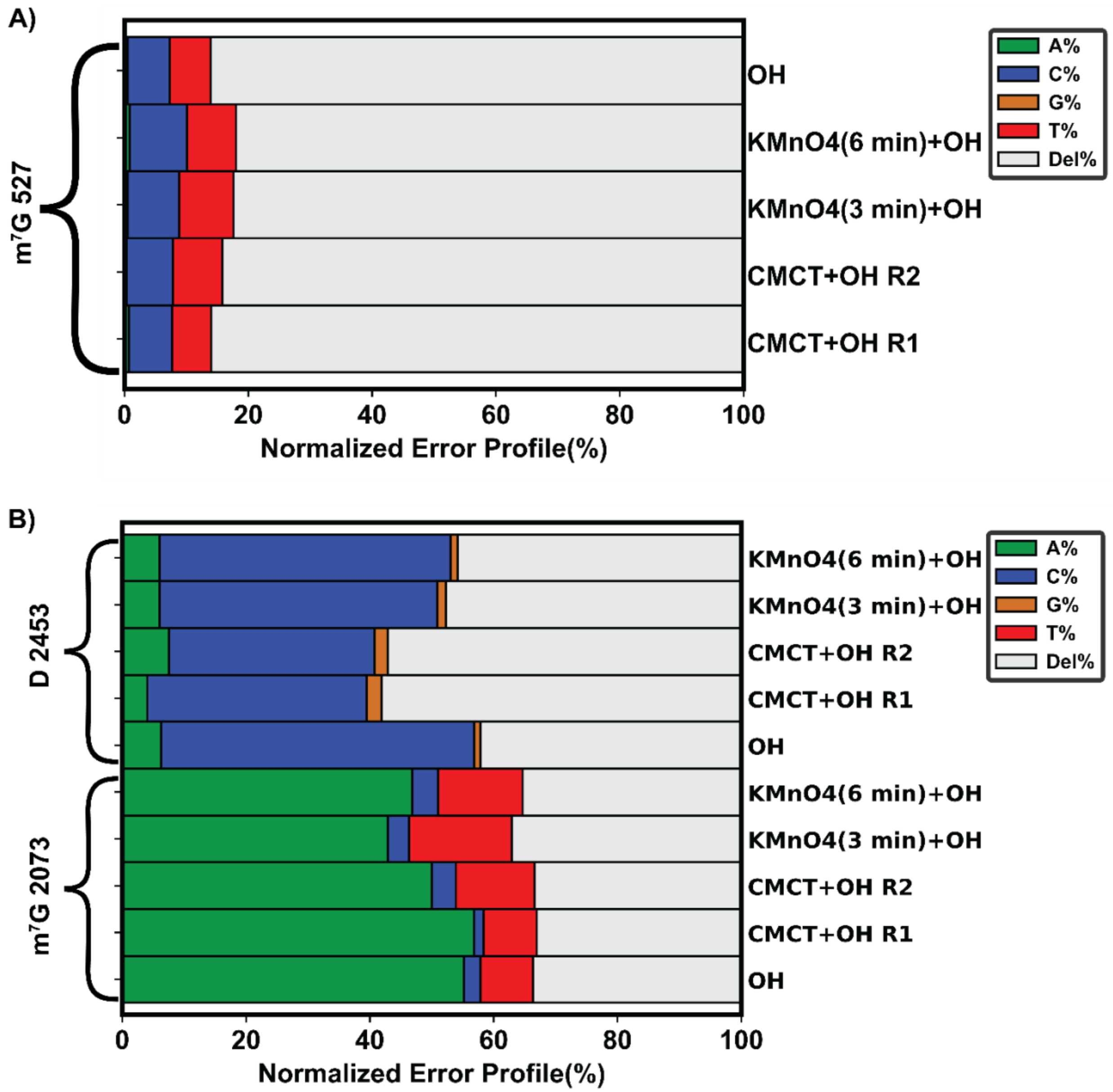

**Figure S2.** The mismatch and deletion profile of $m^7G$ is sequence-context-dependent. (A) Error profiles of $m^7G$ 527 in 16S rRNA of the 30S subunit subjected to the following chemical treatments: $NaHCO_3$; $KMnO_4$ for 3 or 6 minutes, followed by $NaHCO_3$; and CMCT, followed by $NaHCO_3$. (B) Error profiles of $m^7G$ 2073 in 23S rRNA of the 50S subunit subjected to the following chemical treatments: $NaHCO_3$; $KMnO_4$

for 3 or 6 minutes, followed by $NaHCO_3$; and CMCT, followed by $NaHCO_3$. The $m^7G$ 527 modification of 16S rRNA and $m^7G$ 2073 modification of 23S rRNA exhibit the same error profile across different types of chemical treatments. Thus, alkaline treatment induces the same structural change in $m^7G$ regardless of the presence of additional chemical treatments. However, the $m^7G$ 527 modification of 16S rRNA and the $m^7G$ 2073 modification of 23S rRNA induce different reverse transcriptase signatures. The percentages of mismatches and deletions for each modification in this figure were calculated as described in Figure S1. Moreover, the same legend was used for this figure as in Figure S1. In the two panels of this figure, "min" refers to minutes.

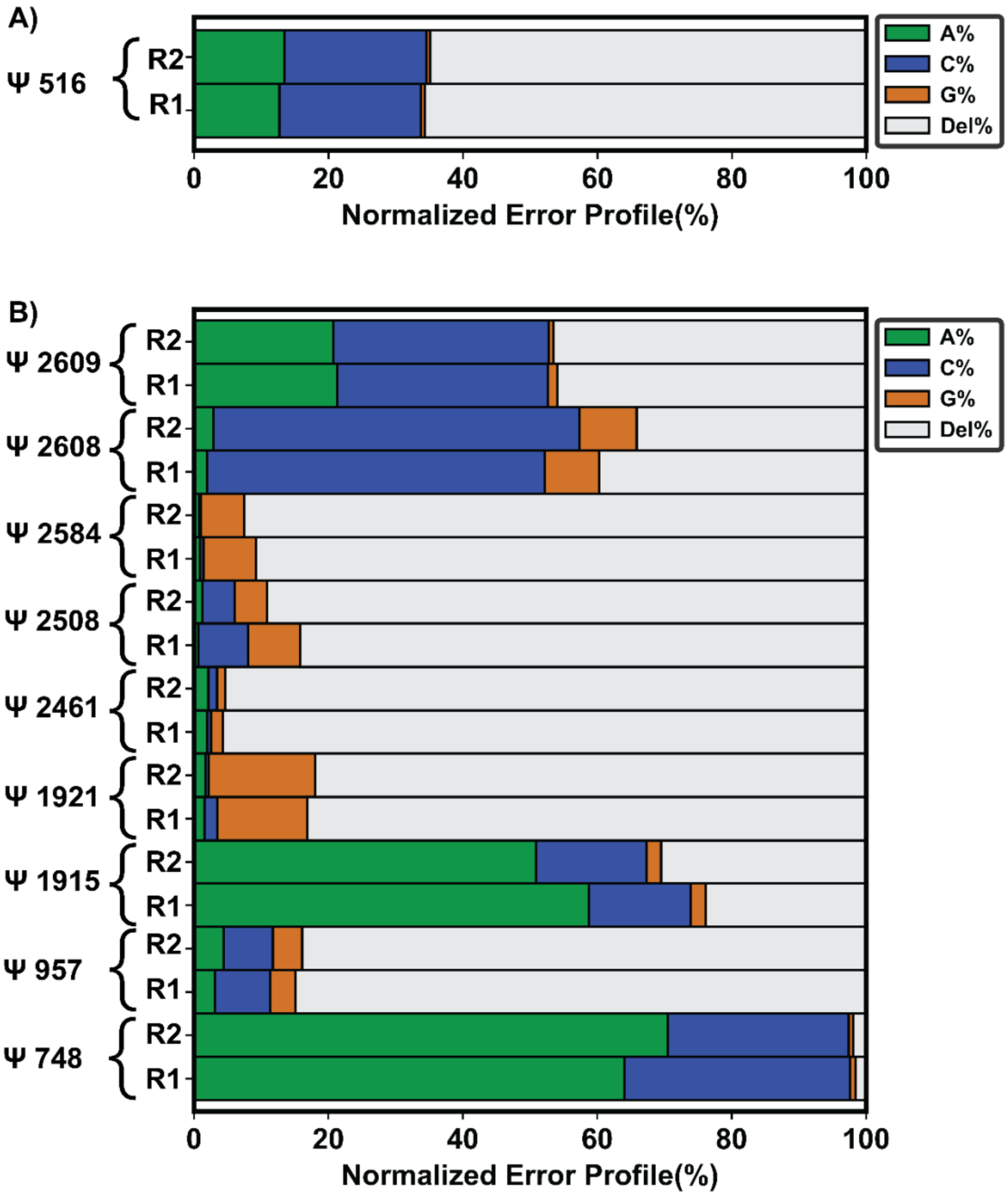

**Figure S3.** The error signatures of reverse transcriptase differ depending on the sequence context of Ψ modifications. In 16S rRNA from the 30S small subunit (A) and the 23S rRNA from the 50S large subunit (B), Ψ modifications detected via CMCT plus $NaHCO_3$ treatment exhibit different error profiles. The sequences where these Ψ modifications are located differ between 16S and 23S rRNA and within the 23S rRNA. Thus, the mismatches and deletions in reverse transcriptase reactions, produced as a consequence of adduct formation between CMCT and Ψ, exhibit a sequence context-dependent profile. In this figure, R1 and R2 present the data obtained from biological replicates 1 and 2. The misincorporation and deletion percentages were calculated as explained in Figure S1. Lastly, the same legend was used as that in Figure S1.

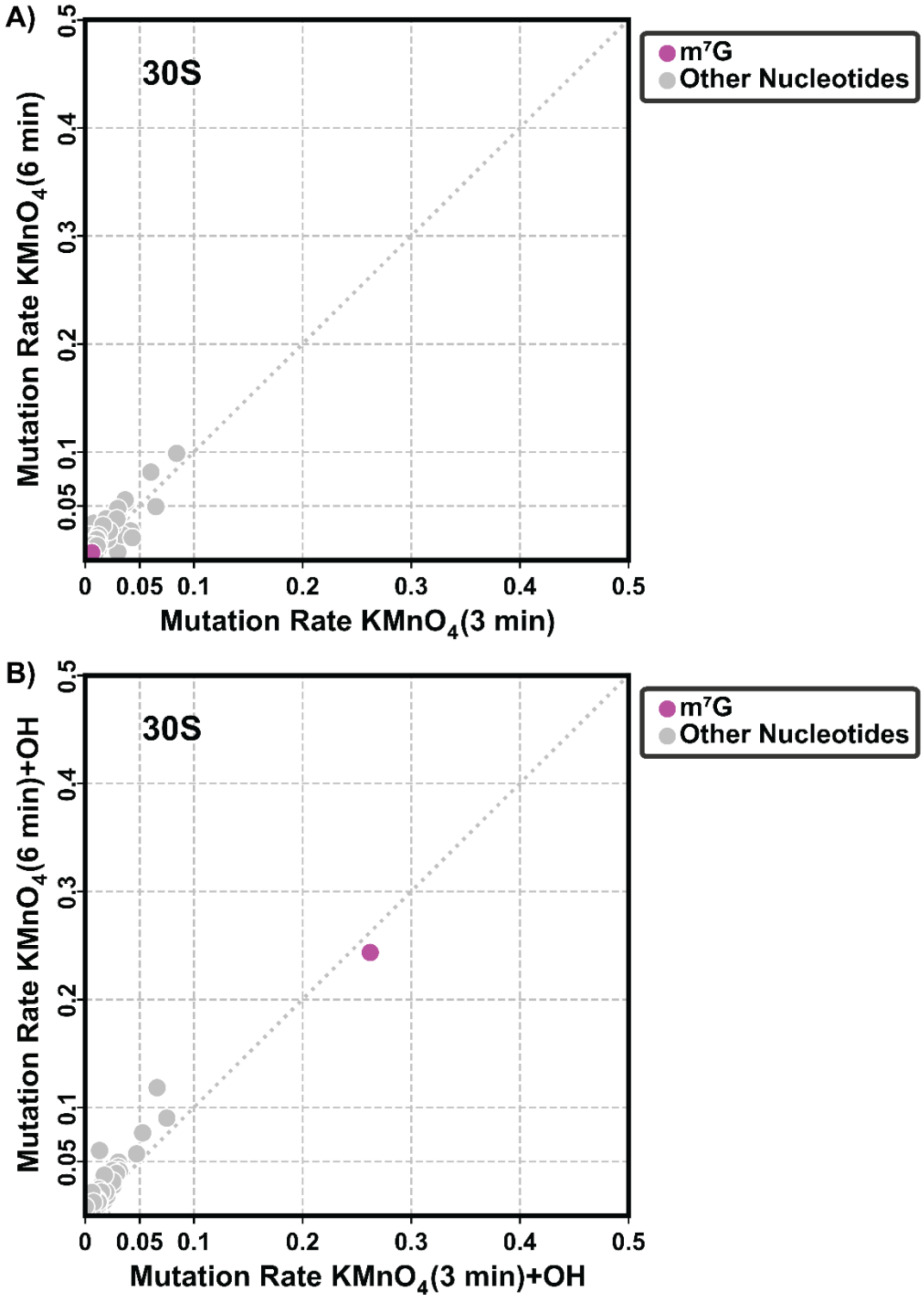
A)
30S
m⁷G
Other Nucleotides
Mutation Rate KMnO4(6 min)
Mutation Rate KMnO4(3 min)
0 0.05 0.1 0.2 0.3 0.4 0.5
B)
30S
m⁷G
Other Nucleotides
Mutation Rate KMnO4(6 min)+OH
Mutation Rate KMnO4(3 min)+OH
0 0.05 0.1 0.2 0.3 0.4 0.5

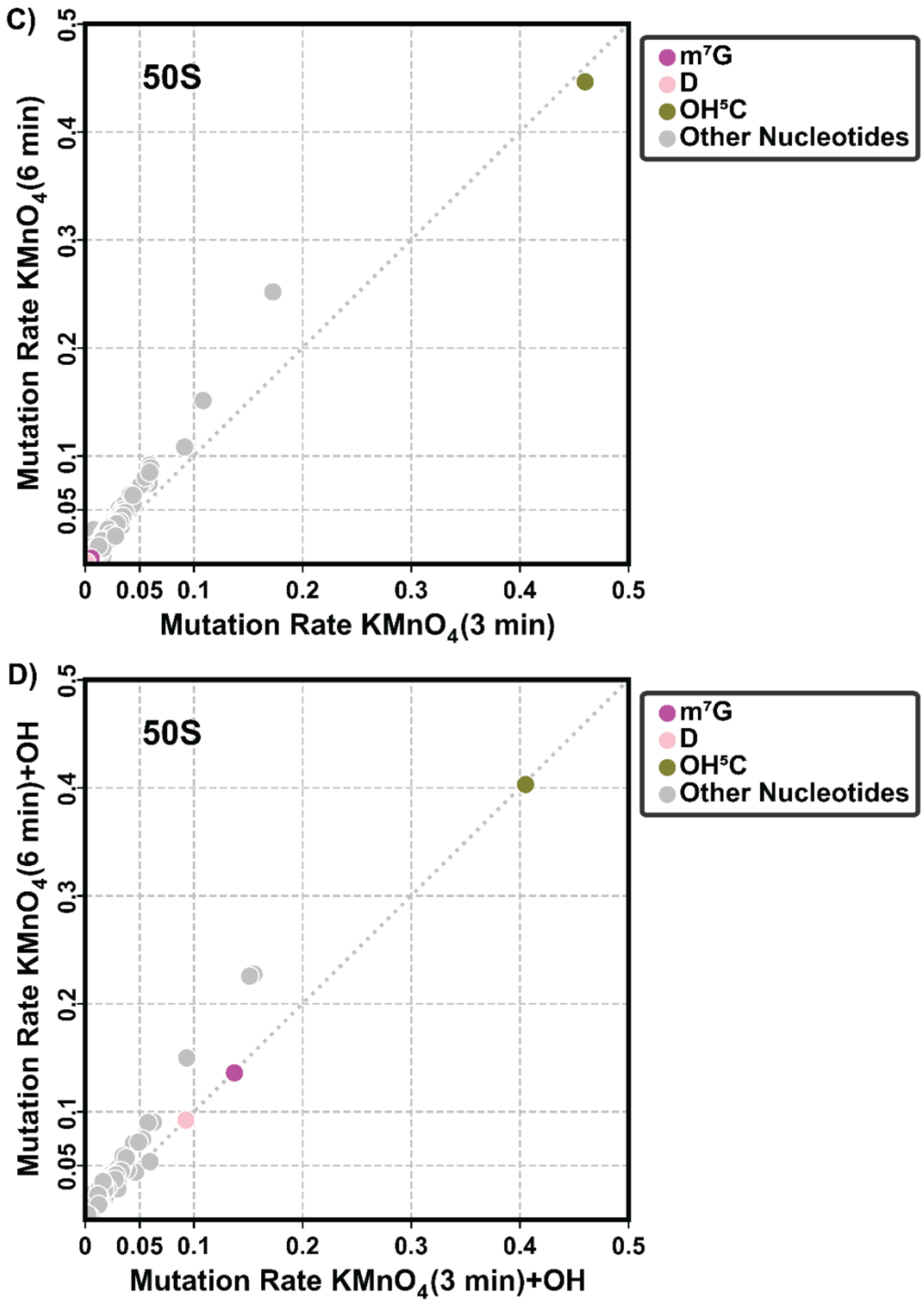

**Figure S4.** D and m[7]G mutation rates are unaffected by $KMnO_4$, whereas they increase in the presence of $NaHCO_3$. A) A comparison of the mutation rates of 16S nucleotides exposed to $KMnO_4$ for 3 (x-axis) and 6 (y-axis) minutes. B) Comparison of the mutation rates for all nucleotides in 16S rRNA samples exposed to $KMnO_4$ for 3 (x-axis) and 6 (y-axis) minutes, followed by exposure to an alkaline treatment. C) A comparison of the mutation rates of all nucleotides in the 23S rRNA from the 50S large subunit subjected to $KMnO_4$ for three (x-axis) or six (y-axis) minutes. $KMnO_4$ does not increase the mutation rates of $m^7G$ and D. On the other hand, as determined in our previous work, it significantly increases the mutation rate of $OH^5C$ compared to the mutation rates of the other nucleotides [32]. D) A comparison of the mutation rates of all nucleotides in the 23S rRNA from the 50S large subunit subjected to $KMnO_4$ for three (x-axis) or six (y-axis) minutes and subsequently exposed to $NaHCO_3$. In the samples subjected to both $KMnO_4$ and $NaHCO_3$, the mutation rates of $m^7G$, D, and $OH^5C$ are higher than those of the other 23S and 16S rRNA nucleotides. A comparison of panels (A), (B), (C), and (D) indicates that the increased mutation rates of residues $m^7G$ and D are a consequence of alkaline treatment. The finding that in the samples treated with both $KMnO_4$ and $NaHCO_3$, the $m^7G$, D, and $OH^5C$ nucleotides have higher mutation rates than the other nucleotides of 23S rRNA demonstrates that the combination of $KMnO_4$ and $NaHCO_3$ could be used to simultaneously detect these three modifications in any RNA molecule. The $m^7G$ modification mutation rates are shown in magenta, D in purple, $OH^5C$ in green, and the other nucleotides of 23S and 16S rRNA in gray. The mutation rates shown in all the panels of this figure were background-corrected as described in Equation 3. In all the panels of this figure, "min" refers to minutes.

**REFERENCES**

[1] Ofengand, J., and Del Campo, M. (2004) Modified Nucleosides of Escherichia coli Ribosomal RNA, *EcoSal Plus 1*.

[2] Okamoto, S., Tamaru, A., Nakajima, C., Nishimura, K., Tanaka, Y., Tokuyama, S., Suzuki, Y., and Ochi, K. (2007) Loss of a conserved 7-methylguanosine modification in 16S rRNA confers low-level streptomycin resistance in bacteria, *Molecular Microbiology 63*, 1096-1106.

[3] Lesnyak, D. V., Osipiuk, J., Skarina, T., Sergiev, P. V., Bogdanov, A. A., Edwards, A., Savchenko, A., Joachimiak, A., and Dontsova, O. A. (2007) Methyltransferase That Modifies Guanine 966 of the 16 S rRNA, *Journal of Biological Chemistry 282*, 5880-5887.

[4] Kimura, S., and Suzuki, T. (2010) Fine-tuning of the ribosomal decoding center by conserved methyl-modifications in the Escherichia coli 16S rRNA, *Nucleic Acids Research 38*, 1341-1352.

[5] Andersen, N. M., and Douthwaite, S. (2006) YebU is a m5C Methyltransferase Specific for 16 S rRNA Nucleotide 1407, *Journal of Molecular Biology 359*, 777-786.

[6] Basturea, G. N., Rudd, K. E., and Deutscher, M. P. (2006) Identification and characterization of RsmE, the founding member of a new RNA base methyltransferase family, *RNA 12*, 426-434.

[7] Basturea, G. N., Dague, D. R., Deutscher, M. P., and Rudd, K. E. (2012) YhiQ Is RsmJ, the Methyltransferase Responsible for Methylation of G1516 in 16S rRNA of E. coli, *Journal of Molecular Biology 415*, 16-21.

[8] Sergiev, P. V., Serebryakova, M. V., Bogdanov, A. A., and Dontsova, O. A. (2008) The ybiN Gene of Escherichia coli Encodes Adenine-N6 Methyltransferase Specific for Modification

of A1618 of 23 S Ribosomal RNA, a Methylated Residue Located Close to the Ribosomal Exit Tunnel, *Journal of Molecular Biology 375*, 291-300.

[9] Sergiev, P. V., Lesnyak, D. V., Bogdanov, A. A., and Dontsova, O. A. (2006) Identification of Escherichia coli m2G methyltransferases: II. The ygjO Gene Encodes a Methyltransferase Specific for G1835 of the 23 S rRNA, *Journal of Molecular Biology 364*, 26-31.

[10] Ero, R., Peil, L., Liiv, A., and Remme, J. (2008) Identification of pseudouridine methyltransferase in Escherichia coli, *RNA 14*, 2223-2233.

[11] Purta, E., O'Connor, M., Bujnicki, J. M., and Douthwaite, S. (2008) YccW is the m5C Methyltransferase Specific for 23S rRNA Nucleotide 1962, *Journal of Molecular Biology 383*, 641-651.

[12] Golovina, A. Y., Dzama, M. M., Osterman, I. A., Sergiev, P. V., Serebryakova, M. V., Bogdanov, A. A., and Dontsova, O. A. (2012) The last rRNA methyltransferase of <i>E. coli</i> revealed: The <i>yhiR</i> gene encodes adenine-N6 methyltransferase specific for modification of A2030 of 23S ribosomal RNA, *RNA 18*, 1725-1734.

[13] Wang, K.-T., Desmolaize, B., Nan, J., Zhang, X.-W., Li, L.-F., Douthwaite, S., and Su, X.-D. (2012) Structure of the bifunctional methyltransferase YcbY (RlmKL) that adds the m 7 G2069 and m 2 G2445 modifications in Escherichia coli 23S rRNA, *Nucleic Acids Research 40*, 5138-5148.

[14] Toubdji, S., Thullier, Q., Kilz, L.-M., Marchand, V., Yuan, Y., Sudol, C., Goyenvalle, C., Jean-Jean, O., Rose, S., Douthwaite, S., Hardy, L., Baharoglu, Z., De Crécy-Lagard, V., Helm, M., Motorin, Y., Hamdane, D., and Brégeon, D. (2024) Exploring a unique class of flavoenzymes: Identification and biochemical characterization of ribosomal RNA dihydrouridine synthase, *Proceedings of the National Academy of Sciences 121*.

[15] Purta, E., O'Connor, M., Bujnicki, J. M., and Douthwaite, S. (2009) YgdE is the 2′-<i>O-</i>ribose methyltransferase RlmM specific for nucleotide C2498 in bacterial 23S rRNA, *Molecular Microbiology 72*, 1147-1158.

[16] Kimura, S., Sakai, Y., Ishiguro, K., and Suzuki, T. (2017) Biogenesis and iron-dependency of ribosomal RNA hydroxylation, *Nucleic Acids Research 45*, 12974-12986.

[17] Toh, S.-M., Xiong, L., Bae, T., and Mankin, A. S. (2008) The methyltransferase YfgB/RlmN is responsible for modification of adenosine 2503 in 23S rRNA, *RNA 14*, 98-106.

[18] Clark, W. C., Evans, M. E., Dominissini, D., Zheng, G., and Pan, T. (2016) tRNA base methylation identification and quantification via high-throughput sequencing, *RNA 22*, 1771-1784.

[19] Dai, Q., Zheng, G., Schwartz, M. H., Clark, W. C., and Pan, T. (2017) Selective Enzymatic Demethylation of <i>N</i><sup>2</sup>,<i>N</i><sup>2</sup>-Dimethylguanosine in RNA and Its Application in High-Throughput tRNA Sequencing, *Angewandte Chemie International Edition 56*, 5017-5020.

[20] Kristen, M., Plehn, J., Marchand, V., Friedland, K., Motorin, Y., Helm, M., and Werner, S. (2020) Manganese Ions Individually Alter the Reverse Transcription Signature of Modified Ribonucleosides, *Genes 11*, 950.

[21] Werner, S., Schmidt, L., Marchand, V., Kemmer, T., Falschlunger, C., Sednev, M. V., Bec, G., Ennifar, E., Höbartner, C., Micura, R., Motorin, Y., Hildebrandt, A., and Helm, M. (2020) Machine learning of reverse transcription signatures of variegated polymerases allows mapping and discrimination of methylated purines in limited transcriptomes, *Nucleic Acids Research 48*, 3734-3746.

[22] Araujo Tavares, R. D. C., Mahadeshwar, G., Wan, H., and Pyle, A. M. (2023) MRT-ModSeq – Rapid detection of RNA modifications with MarathonRT, Cold Spring Harbor Laboratory.

[23] Koculi, E., and Cho, S. S. (2022) RNA Post-Transcriptional Modifications in Two Large Subunit Intermediates Populated in <i>E. coli</i> Cells Expressing Helicase Inactive R331A DbpA, *Biochemistry 61*, 833-842.

[24] Narayan, G., Gracia Mazuca, L. A., Cho, S. S., Mohl, J. E., and Koculi, E. (2023) RNA Post-transcriptional Modifications of an Early-Stage Large-Subunit Ribosomal Intermediate, *Biochemistry 62*, 2908-2915.

[25] Finet, O., Yague-Sanz, C., Krüger, L. K., Tran, P., Migeot, V., Louski, M., Nevers, A., Rougemaille, M., Sun, J., Ernst, F. G. M., Wacheul, L., Wery, M., Morillon, A., Dedon, P., Lafontaine, D. L. J., and Hermand, D. (2022) Transcription-wide mapping of dihydrouridine reveals that mRNA dihydrouridylation is required for meiotic chromosome segregation, *Molecular Cell 82*, 404-419.e409.

[26] Draycott, A. S., Schaening-Burgos, C., Rojas-Duran, M. F., Wilson, L., Schärfen, L., Neugebauer, K. M., Nachtergaele, S., and Gilbert, W. V. (2022) Transcriptome-wide mapping reveals a diverse dihydrouridine landscape including mRNA, *PLOS Biology 20*, e3001622.

[27] Marchand, V., Bourguignon-Igel, V., Helm, M., and Motorin, Y. (2021) Chapter Two - Mapping of 7-methylguanosine (m7G), 3-methylcytidine (m3C), dihydrouridine (D) and 5-hydroxycytidine (ho5C) RNA modifications by AlkAniline-Seq, In *Methods in Enzymology* (Jackman, J. E., Ed.), pp 25-47, Academic Press.

[28] Marchand, V., Ayadi, L., Bourguignon-Igel, V., Helm, M., and Motorin, Y. (2021) AlkAniline-Seq: A Highly Sensitive and Specific Method for Simultaneous Mapping of 7-Methyl-guanosine (m7G) and 3-Methyl-cytosine (m3C) in RNAs by High-Throughput Sequencing, pp 77-95, Springer US.

[29] Enroth, C., Poulsen, L. D., Iversen, S., Kirpekar, F., Albrechtsen, A., and Vinther, J. (2019) Detection of internal N7-methylguanosine (m7G) RNA modifications by mutational profiling sequencing, *Nucleic Acids Research 47*, e126-e126.

[30] Zhang, L.-S., Liu, C., Ma, H., Dai, Q., Sun, H.-L., Luo, G., Zhang, Z., Zhang, L., Hu, L., Dong, X., and He, C. (2019) Transcriptome-wide Mapping of Internal N7-Methylguanosine Methylome in Mammalian mRNA, *Molecular Cell 74*, 1304-1316.e1308.

[31] Zhang, L.-S., Ju, C.-W., Liu, C., Wei, J., Dai, Q., Chen, L., Ye, C., and He, C. (2022) m<sup>7</sup>G-quant-seq: Quantitative Detection of RNA Internal <i>N</i><sup>7</sup>-Methylguanosine, *ACS Chemical Biology 17*, 3306-3312.

[32] Koculi, E., and Cho, S. S. (2022) RNA Post-Transcriptional Modifications in Two Large Subunit Intermediates Populated in E. coli Cells Expressing Helicase Inactive R331A DbpA, *Biochemistry 61*, 833-842.
